# *In vivo* dissection of a clustered-CTCF domain boundary reveals developmental principles of regulatory insulation

**DOI:** 10.1101/2021.04.14.439779

**Authors:** Chiara Anania, Rafael D. Acemel, Johanna Jedamzick, Adriano Bolondi, Giulia Cova, Norbert Brieske, Ralf Kühn, Lars Wittler, Francisca M. Real, Darío G. Lupiáñez

## Abstract

Vertebrate genomes organize into topologically associating domains (TADs), delimited by boundaries that insulate regulatory elements from non-target genes. However, how boundary function is established is not well understood. Here, we combine genome-wide analyses and transgenic mouse assays to dissect the regulatory logic of clustered-CTCF boundaries *in vivo*, interrogating their function at multiple levels: chromatin interactions, transcription and phenotypes. Individual CTCF binding sites (CBS) deletions revealed that the characteristics of specific sites can outweigh other factors like CBS number and orientation. Combined deletions demonstrated that CBS cooperate redundantly and provide boundary robustness. We show that divergent CBS signatures are not strictly required for effective insulation and that chromatin loops formed by non-convergently oriented sites could be mediated by a loop interference mechanism. Further, we observe that insulation strength constitutes a quantitative modulator of gene expression and phenotypes. Our results highlight the modular nature of boundaries and their control over developmental processes.

## INTRODUCTION

The development of complex organisms relies on intricate gene expression patterns that are essential for the proper differentiation of tissues and cell types. In vertebrates, a major means of achieving transcriptional control is through the action of distal regulatory elements, such as enhancers (Long et al., 2016). To elicit a precise transcriptional response, regulatory elements are required to come into physical proximity with their target gene promoters. Such functional interaction is mediated by the 3D folding of the chromatin, which facilitates the regulatory interplay between regions otherwise distant on a linear genome. In recent years, substantial efforts have been directed towards a better characterization of the molecular mechanisms that drive 3D chromatin folding and on their influence on developmental processes. The emergence of high-throughput conformation capture methods (Hi-C) has allowed a detailed investigation of nuclear interactions (Lieberman-Aiden et al., 2009; de Wit and de Laat, 2012), revealing that vertebrate genomes organize into topologically associating domains (TADs) (Dixon et al., 2012; Nora et al., 2012). TADs represent megabase-sized regions containing loci with increased interaction frequencies and, perhaps more importantly, they often constitute functional domains in which regulatory elements and their cognate genes are framed (Shen et al., 2012; Symmons et al., 2014). TADs are separated by boundaries, which are genomic regions with insulating properties that limit the regulatory crosstalk between adjacent regulatory domains. TAD boundaries represent an important regulatory hallmark along the genome, as their disruption has been linked to human disease, including congenital malformations and cancer (Flavahan et al., 2016; Hnisz et al., 2016; Lupiáñez et al., 2015; Spielmann et al., 2018).

Genomic analyses of TAD boundaries regions identified the transcriptional repressor CCCTC-binding factor (CTCF) as a key player in 3D chromatin organization (Ong and Corces, 2014). CTCF is found at the majority of TAD boundaries, where its binding mediates the insulation properties of these regions (Dixon et al., 2012). Consistent with this notion, CTCF depletion leads to a genome-wide disappearance of TADs (Nora et al., 2017), thus providing mechanistic insights into CTCF genome binding properties. The genomic distribution of CTCF is particularly influenced by the orientation of its DNA binding motif. The formation of chromatin loops, often associated with TAD boundaries, preferentially occurs between pairs of CTCF binding sites (CBS) displaying convergent motif orientations (Rao et al., 2014). Accordingly, the inversion of CTCF motifs can redirect chromatin loops (Guo et al., 2015; de Wit et al., 2015). At TAD boundaries, the clustering of CBS with divergent orientation is a recurrent molecular signature that has been conserved through vertebrate evolution (Gómez-Marín et al., 2015). From a mechanistic perspective, the orientation bias of CTCF can be explained by the loop extrusion model (Fudenberg et al., 2016; Sanborn et al., 2015). This model proposes that the cohesin complex extrudes the chromatin fiber until reaching a CTCF site bound in an opposing orientation, but continuing when CTCF is oriented otherwise. As occurs for CTCF, the depletion of the cohesin complex leads to a global loss of TAD insulation (Rao et al., 2017; Schwarzer et al., 2017; Wutz et al., 2017), revealing a tight regulatory interplay between these two architectural factors.

Based on this experimental evidence, it is assumed that boundary elements and CTCF are fundamental players for the spatial organization of genomes. However, the degree to which TAD boundaries influence developmental gene expression remains highly controversial. While alterations of TAD boundaries at particular loci can lead to developmental phenotypes or cancer (Flavahan et al., 2016; Franke et al., 2016; Hnisz et al., 2016; Lupiáñez et al., 2015), it only caused moderate transcriptional changes in other genomic regions (Despang et al., 2019; Paliou et al., 2019; Williamson et al., 2019). In addition, the global disruption of TADs via CTCF or cohesin depletion in cultured cells only results in limited changes in gene expression (Nora et al., 2017; Rao et al., 2017). Furthermore, single-cell Hi-C (Flyamer et al., 2017; Stevens et al., 2017) or super-resolution microscopy studies (Bintu et al., 2018) have revealed that individual cells can display chromatin conformations that, in some instances, ignore the TAD boundaries detected in aggregated or bulk datasets. Such contradictory results demonstrate the need for a comprehensive dissection of boundary elements in developmental settings.

Here, we combine genome-wide analyses with the systematic dissection of a TAD boundary in transgenic mice to investigate the molecular principles of boundary function *in vivo*. Using the *Epha4-Pax3* (EP) boundary region as a testbed for experimental validations, we generated a collection of 13 mouse homozygous alleles carrying individual or combined CBS deletions that covers a broad range of regulatory configurations. We subsequently combined capture Hi-C (cHi-C), gene expression and phenotypical analyses to quantify the functional consequences of these boundary perturbations at several levels: ectopic chromatin interactions, gene misexpression and aberrant limb morphologies. We discover that functional characteristics of specific CBS are major determinants of boundary insulation, outweighing other relevant parameters such as the number or the orientation of CBS. By performing combined deletions, we reveal that CBS cooperate to achieve precise levels of insulation. Nevertheless, they are also partially redundant, a property that confers robustness to boundary regions. Further, we show that a divergent CBS signature is not a strict requisite for efficient boundary function and that CBS with a strong insulator function can also establish chromatin loops in non-convergent orientations, for which we suggest a mechanism of loop interference. Furthermore, we observe that insulation strength influences gene expression and phenotypes, by quantitatively modulating the degree of regulatory interactions across adjacent TADs. Our results reveal fundamental principles of boundary elements and delineate a tight interplay between genomic sequence, 3D chromatin structure and developmental function.

## RESULTS

### A genetic setup to investigate boundary function in vivo

By studying a series of deletions in transgenic mice, we previously demonstrated that a 150 Kilobases (Kb) region, marked as a boundary region across multiple tissues and cell types, is sufficient to segregate the regulatory activities of the *Epha4* and *Pax3* TADs (Lupiáñez et al., 2015) **(Supp. Fig. 1 and 2)**. The *DelB* background carries a deletion that removes a portion of the *Epha4* TAD, including the gene itself, as well as the boundary region that separates this domain from the adjacent *Pax3* TAD. This deletion results in the ectopic interaction between the *Epha4* limb enhancers and the *Pax3* gene, which causes the misexpression of *Pax3* in developing limbs and leads to the shortening of index and thumb fingers (brachydactyly) in mice and also in human patients with equivalent deletions. In contrast, the *DelBs* background carries a similar deletion but not affecting the EP boundary region, which maintains the regulatory partition between the *Epha4* and *Pax3* TADs and confines the *Epha4* limb-specific enhancers within their own regulatory domain **(Fig. 1A, Supp. Fig. 1)**. Thereby, the genomic configuration of the *DelBs* background provides a simple, but informative, functional readout to investigate boundary function *in vivo*. By performing deletions on the genomic components of the EP boundary, we can quantify the consequences of boundary disruption on a single target gene that is reactive to ectopic enhancers and can induce developmental defects. This genomic setup allows us to estimate boundary function at multiple levels: inter-TAD chromatin interactions, gene misexpression and disease-related phenotypes.

**Figure 1.**
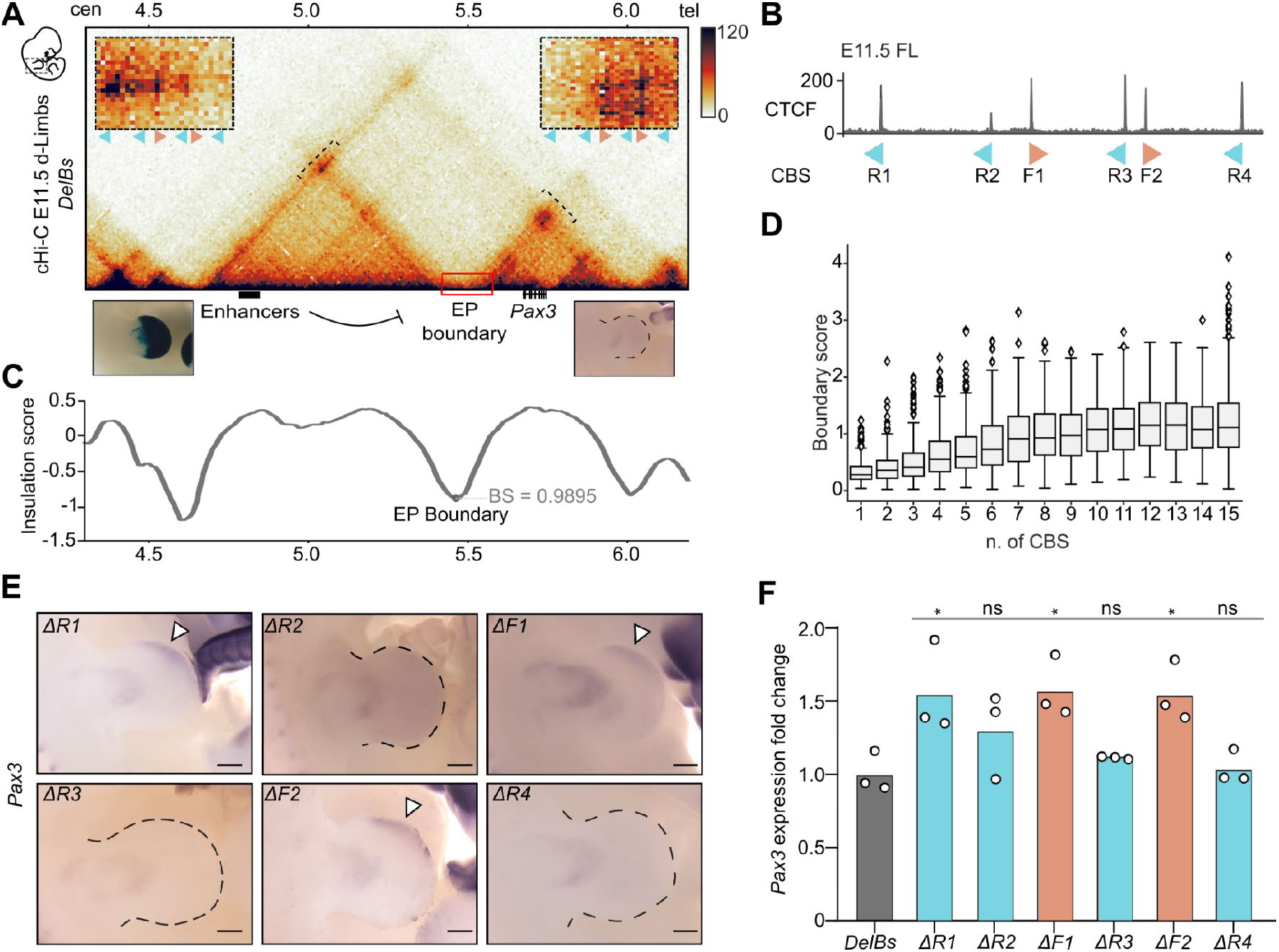
Impact of individual CBS deletions on boundary function. **A.** c-HiC maps from E11.5 distal limbs from *DelBs* mutants at 10 kb resolution. Data mapped on custom genome containing the *DelBs* deletion. Red rectangle marks the EP boundary region. Insets represent a magnification (5kb resolution) of the centromeric (left) and telomeric (right) loops highlighted by brackets on the map. Arrowheads represent reverse (light blue) and forward (orange) oriented CBS. Below, Lac-Z staining (left) and WISH (right) of E11.5 mouse forelimbs show activation pattern of *Epha4* enhancers and *Pax3* expression respectively. **B.** CTCF ChIP-seq track in E11.5 mouse forelimbs (Andrey et al., 2017). Schematic shows CBS orientation. **C.** Insulation score values. Gray dot represents the local minima of the insulation score at EP boundary, also measured as boundary strength (BS). **D.** Relationship between Boundary Scores (BS) and the number of CBS (data from (Bonev et al., 2017)). The boxes in the boxplots indicate the median and the first and third quartiles (Q1 and Q3). Whiskers extend 1.5 times the interquartile range below and above Q1 and QR respectively. **E.** WISH shows *Pax3* expression in E11.5 forelimbs from CBS mutants. Note *Pax3* misexpression on the distal anterior region in *ΔR1, ΔF1 and ΔF2* mutants (white arrowheads). Scale bar 250 μm. **F.** *Pax3* qPCR analysis in E11.5 limb buds from CBS mutants. Bars represent the mean and white dots represent individual replicates. Values normalized against *DelBs* mutant (ΔΔCt). (* p-value ≤ 0.05; ns: non-significant).

To explore the genomic features of the EP boundary *in vivo*, we examined ChIP-seq datasets on developing limbs (Rodríguez-Carballo et al., 2017). This analysis revealed the presence of six clustered CBS at the EP boundary region (**Fig. 1A, 1B; Supp Fig 2**). CTCF motif analyses confirmed the divergent orientation of these sites, a typical signature of TAD boundaries, with four CBS in a reverse (R) and two in a forward orientation (F). Overall, the profile of CTCF binding at the EP boundary is conserved across tissues, despite quantitative variations at individual sites (**Supp. Fig. 2**) (Bonev et al., 2017; Rodríguez-Carballo et al., 2017). Other prominent features associated with boundary regions, such as active transcription or housekeeping genes were not found in the region (**Supp. Fig. 3**). High-resolution Capture Hi-C (cHi-C) data from *DelBs* E11.5 distal limbs revealed that the EP boundary region establishes chromatin loops with the immediately centromeric and telomeric TAD boundaries (Bianco et al., 2018). Consistent with the convergent CTCF orientation bias (Guo et al., 2015; Rao et al., 2014; de Wit et al., 2015), two chromatin loops connect the two forward-oriented CBS (F1 and F2) with the telomeric boundary of the *Pax3* TAD **(Fig. 1A and B)**. Similarly, chromatin loops are also established between the centromeric boundary of the *Epha4* TAD and the reverse-oriented CBS R1, R2 and R3 (**Fig. 1A**). However, the close genomic distances between R2 and F1 and between R3 and F2 precludes the unambiguous assignment of chromatin loops to specific sites. Intriguingly, only R1, F1 and F2 are specifically bound by the cohesin complex (Andrey et al., 2017) (**Supp. Fig. 3**), an essential component for the formation of chromatin loops (Rao et al., 2017; Schwarzer et al., 2017; Wutz et al., 2017). Overall, these results delineate the EP element as a prototypical boundary region with insulating properties that are likely encoded and controlled by CBS.

### The characteristics of individual CTCF binding sites are major determinants of boundary function

Boundary regions are predominantly composed of CBS clusters (Kentepozidou et al., 2020), suggesting that the number of sites might be relevant for their function. We explored this hypothesis by calculating boundary scores (Crane et al., 2015), a proxy for insulator function, on available ultra-high resolution Hi-C maps (Bonev et al., 2017). Specifically, we employed this parameter to identify boundary regions, which we further categorized according to the CBS number, identified by CTCF ChIP-seq experiments. Overall, we observe that boundary scores increase monotonically with the number of clustered CBS, reaching a stabilization at 10 CBS (**Fig. 1D**). According to this distribution, the EP boundary would fall within a range where its function might be sensitive to alterations on the CBS number. To test this, we took advantage of a mouse homozygous embryonic stem cell (mESC) line for the *DelBs* background (Bianco et al., 2018). Using CRISPR/Cas9 technology, we systematically targeted the *DelBs* mESCs to generate individual homozygous deletions for each of the six CBS that constitute the EP boundary region. For each deletion, we selected clones that display a successful disruption of the CTCF binding motif in both alleles (**Supp. Fig 4**). ChIP-seq experiments revealed that, in all cases, the disruption of the binding motif was sufficient to completely abolish CTCF recruitment at the targeted site (**Supp. Fig. 5**). Subsequently, we employed tetraploid complementation assays to generate mutant embryos and measure the functional consequences of these individual deletions *in vivo* (Artus and Hadjantonakis, 2011; Kraft et al., 2015).

Whole mount in situ hybridization (WISH) on E11.5 mutant embryos revealed that the insulation function of the EP boundary could be sensitive to individual perturbations of CBS (**Fig. 1E**). However, this effect was not observed for each CBS deletion, but only for those previously associated with RAD21 binding (R1, F1 and F2) (**Fig. 1A; Supp. Fig. 3**). For *ΔR1, ΔF1* and *ΔF2* mutants, the alteration of boundary function is evidenced by the ectopic expression of *Pax3* on a reduced area of the anterior limb. The expression domains of *Pax3* in other tissues remained unaltered (**Supp. Fig. 6**), thus confirming that changes of its endogenous pattern of expression are restricted to the limb bud. The disruption of the other CBS of the boundary (*ΔR2, ΔR3* and *ΔR4* mutants) did not result in observable changes in *Pax3* expression, demonstrating that the EP boundary can also preserve its function despite a reduction in the total number of CBS.

To quantify the levels of *Pax3* misexpression, we performed quantitative PCR (qPCR) experiments in developing forelimbs at E11.5. We observe a modest, but significant, upregulation in *ΔR1, ΔF1* and *ΔF2* mutants, compared to *DelBs* control animals (1.5-fold upregulation) (**Fig 1F**). No significant differences were detected in the other mutants, in agreement with the results obtained from WISH experiments. Importantly, we observe that the functionality of individual CBS is not strictly correlated with CTCF occupancy, as the deletion of R3, which displays the highest levels of CTCF binding among the cluster (**Supp. Fig. 3**), does not result in measurable gene expression changes (**Fig. 1F**). Overall, these results suggest that, while the CBS number influences insulation, the individual characteristics of specific sites constitute major determinants of boundary function.

### CTCF binding sites cooperate redundantly to provide insulator robustness

The perturbed boundary insulation observed across several mutants (**Fig. 1E and F**), as well as the correlative increase of boundary strength and CBS number (**Fig. 1D**), may suggest functional cooperation between sites. To further explore this hypothesis, we retargeted our *ΔR1* mESC line to generate double knockout mutants of CBS for which individual deletions led to reduced insulation. We chose CBS combinations with either different (R1 and F2 for *ΔR1+F2*) or identical orientations (F1 and F2 for *ΔF-all*) (**Fig. 2A**). Both combined CBS deletions led to an expansion of *Pax3* misexpression towards the posterior region of the limb, suggesting that the EP boundary is further compromised in the double mutants. These results are consistent with our previous observations, supporting an important role of CBS F1, F2 and R1 on EP boundary function (**Fig. 1B**). Next, we sought to determine the nature of CBS cooperation by performing qPCR analyses in developing limbs (**Fig. 2B**). The increased *Pax3* misexpression exceeded the summed expression levels from the corresponding individual deletions, for both mutants. In particular, *ΔR1+F2* mutants display an increase in *Pax3* misexpression of up to 6.3-fold compared to *DelBs* controls, contrasting with the 1.8-fold and 1.5-fold observed in *ΔR1* and *ΔF2* mutants. For *ΔF-all* mutants, this effect is even more prominent, as denoted by a 9.1-fold increase of *Pax3* misexpression compared to the 1.8-fold of *ΔF1* and 1.6-fold increase of *ΔF2* mutants. The negative epistatic effects of the combined mutations indicate that the different CBS of the boundary display partially redundant functions, compensating for the absence of each other.

**Figure 2.**
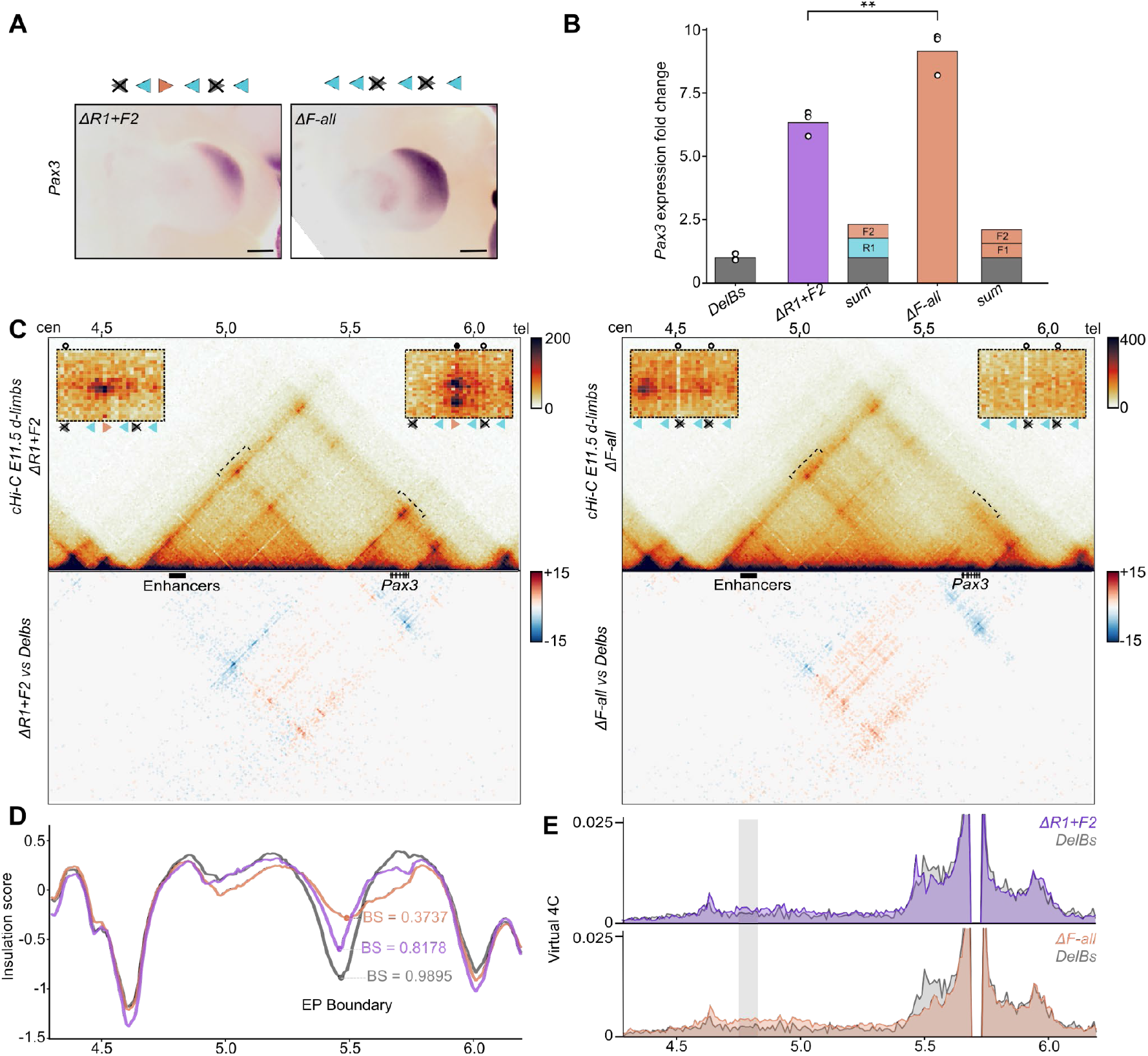
Impact of combined CBS deletions on boundary function. **A.** WISH shows *Pax3* expression in E11.5 forelimbs from CBS mutants. Arrowheads represent reverse (light blue) and forward (orange) oriented CBS. Crosses indicate deleted CBS. Note increased *Pax3* misexpression towards the posterior regions of the limbs. Scale bar: 250μm. **B.** *Pax3* qPCR analysis in E11.5 limb buds from CBS mutants. Bars represent the mean and white dots represent individual replicates. Values normalized against *DelBs* mutant (ΔΔCt). (* p-value ≤ 0.05; ns: non-significant). **C.** c-HiC maps from E11.5 mutant distal limbs at 10 kb resolution (top). Data mapped on custom genome containing the *DelBs* deletion. Insets represent a magnification (5kb resolution) of the centromeric (left) and telomeric (right) loops highlighted by brackets on the map. Gained or lost chromatin loops represented by full or empty dots, respectively. Subtraction maps (bottom) showing gain (red) or loss (blue) of interactions in mutants compared to *DelBs*. **D.** Insulation score values. Lines represent indicated mutants. Dots represent the local minima of the insulation score at EP boundary for each mutant, also measured as boundary score (BS). **E.** Virtual 4C profiles with *Pax3* promoter as a viewpoint for the genomic region displayed in panel **C**. Light gray rectangle highlights *Epha4* enhancer region. Note increased interactions between *Pax3* promoter and *Epha4* enhancer in *ΔR1+F2* and *ΔF-all* (purple and orange) compared to *DelBs* mutants (gray).

To gain further insights on the molecular mechanisms of CBS cooperation, we generated cHi-C maps of the *Epha4-Pax3* locus on distal limbs from mutant E11.5 embryos (**Fig. 2C; Supp Fig. 7**). Interaction maps from *ΔR1+F2* embryos denoted a clear partition between the *EphaA4* and *Pax3*, analogous to *DelBs* control mutants (**Fig. 2C**). However, subtraction maps against *DelBs* controls showed a decrease in the intra-TAD interactions for both the *Epha4* and *Pax3* TADs, concomitant with an increase in the interactions between these two domains. Accordingly, the boundary scores of the EP boundary in *ΔR1+F2* mutants decreased 17% compared to *DelBs*, reflecting a weakened insulating capacity (**Fig. 2D**). Virtual 4C-profiles demonstrated that the perturbation of the EP boundary leads to increased chromatin interactions between the *Pax3* promoter and the *Epha4* limb enhancers (**Fig. 2E**), consistent with the upregulation of *Pax3*. Additionally, the combined disruption of R1 and F2 also induced relevant structural changes in the 3D organization of the locus. Two of the chromatin loops that connect the EP boundary and the telomeric boundary were abolished, due to the deletion of the F2 CBS anchor (**Fig. 2C; Supp. Fig. 8**). Consequently, the adjacent chromatin loop exhibited a compensatory effect, with a notable increase in the interactions mediated by the F1 anchor. At the centromeric site, the deletion of R1 causes the relocation of the loop anchor towards an adjacent region containing the reverse CBS (R2) and the only remaining forward CBS (F1). While the loop extrusion model would indeed predict a stabilization at a reverse CBS (Fudenberg et al., 2016; Sanborn et al., 2015), the short genomic distance between R2 and F1 does not allow us to assign the anchor point for this loop unambiguously. Moreover, we observed the presence of contacts at R3 and R4, suggesting that these sites are also functionally redundant.

Then, we examined cHi-C maps from *ΔF-all* mutants, from which we had detected a more pronounced *Pax3* misexpression (**Fig. 2B**; 1.45-fold compared to the *ΔR1+F2* mutant). Interaction maps revealed a partial fusion of the *Epha4* and *Pax3* domains (**Fig. 2C**), accompanied by a notable decrease of the boundary score at the EP boundary (**Fig. 2D;** boundary score decreased by 62%). Virtual 4C profiles confirmed the increase in interactions between *Pax3* and the *Epha4* enhancers in *ΔF-all* compared to *ΔR1+F2* mutants, in agreement with the more pronounced *Pax3* upregulation (**Fig. 2E**). As for *ΔR1+F2* mutants, we also observed differences in *ΔF-all* interaction maps that can be attributed to the perturbation of specific CBS (**Fig. 2C; Supp. Fig. 8**). In particular, the deletion of all CBS with forward orientation abolishes all chromatin loops that connect with the telomeric *Pax3* boundary. Towards the centromeric side, R1 maintains its chromatin loop with the centromeric *Epha4* boundary. However, other chromatin loops are still discernible and anchored by the R3 and R4 sites, confirming previous indications that these sites perform distinct yet partially overlapping functions. Overall, these results demonstrate that, in the context of a boundary, CBS cooperate but also can partially compensate for the absence of each other, therefore conferring functional robustness to boundary regions.

### Formation of chromatin loops by non-convergent CTCF binding sites through loop interference

As previously mentioned, chromatin loops are predominantly anchored by CBS pairs displaying convergent motif orientation, with both anchor sites oriented inwards with respect to the loop (Rao et al., 2014; de Wit et al., 2015). Intriguingly, we observed that the combined deletion of the forward F1 and F2 sites (*ΔF-all*) not only disrupts the telomeric loops, but also impacts the centromeric one, an effect that is visible in the subtraction maps (**Fig. 2C**). This effect is noticeable at the R2/F1 site, which was associated with a centromeric chromatin loop in the *DelBs* background (**Fig. 1A**). The disruption of this loop in the *ΔF-all* mutant demonstrates that its main anchor point was not the R2 but the F1 site (**Supp. Fig 8**). ChIP-seq analyses in limbs showed differences in CTCF binding between these two sites that might partially explain the preferential anchoring (**Fig. 1B**). Nevertheless, these results also suggest that the F1 CBS can form chromatin loops in a non-convergent orientation. A plausible mechanistic explanation is described by the loop extrusion model, which predicts that existing chromatin loops could create steric impediments that might prevent additional cohesin complexes from sliding through anchor sites (Fudenberg et al., 2016; Sanborn et al., 2015). This effect would stabilize these additional cohesin complexes, resulting in the establishment of simultaneous and paired non-convergent and convergent loops, which would manifest as a similar structure as observed in our cHi-C maps.

We searched for further biological indications of this mechanism by analyzing ultra-high resolution Hi-C datasets (Bonev et al., 2017). First, we identified loop anchors and classified them according to the motif orientation of their CBS and the loops they form. Namely, loop anchors were split in convergent-only (if they always contain CBS oriented in the same direction as their anchored loops), non-convergent (if they anchor at least one loop in a direction for which they lack a directional CBS) and no-CTCF (if they do not contain any CBS). Consistent with previous reports, most loop anchors belong to the convergent-only category (Rao et al., 2014; de Wit et al., 2015). However, 7.6% of them were classified as non-convergent. Then, we explored whether these non-convergent loops could be explained by the fact that the non-convergent anchor simultaneously establishes a convergent loop in the opposite direction (**Fig. 3A**). We calculated the frequency of anchors involved in loops in both directions in each of the three different categories and discovered that, while only 5% of convergent-only or no-CTCF anchors participate in bidirectional loops, this percentage increases significantly up to 45% for non-convergent anchors (**Fig. 3B**; χ2 p-val < 10^-225^). Therefore, non-convergent anchors establish loops in both directions more often than convergent-only and no-CTCF anchors. To gain further insights into the mechanisms that establish convergent/non-convergent pairs of loops, we calculated the strength of each corresponding paired loop (Flyamer et al., 2017). This analysis revealed that the convergent loops linked to a non-convergent loop are significantly stronger than their non-convergent mates (**Fig. 3C and D**; Mann-Whitney U p-value=6×10^-6^). Next, we explore if convergent loops paired to non-convergent loops are particularly strong in comparison to other types of convergent loops. This analysis revealed that the strength of these convergent loops is similar to other unpaired convergent loops across the genome (single-sided convergent category; **Supp. Fig. 9**). However, paired convergent/non-convergent loops appear to be mechanistically different from unpaired loops, as they are more often associated with TAD corners (χ2 p-val<3.5×10^-6^, **Supp. Fig. 9C**) and therefore connect anchor points that are located farther away in the linear genome (Mann-Whitney U p-value <4.8×10^-8^, **Supp. Fig. 9D**). A comparison against pairs of convergent/convergent loops, which are similarly associated with TAD-corners (category double-sided convergent in **Supp. Fig. 9B**), revealed that the convergent loops in convergent/non-convergent pairs are on average stronger (Mann-Whitney U p-val = 7×10^-5^). Importantly, this type of convergent/non-convergent loops can be observed at relevant developmental loci, such as the *Osr1, Ebf1* and *Has2* (**Supp. Fig. 10**). Overall, our analyses suggest that a considerable number of non-convergent loops could be mechanistically explained by the presence of a stronger and convergent chromatin loop in the opposite orientation and anchored by the same CBS.

**Figure 3.**
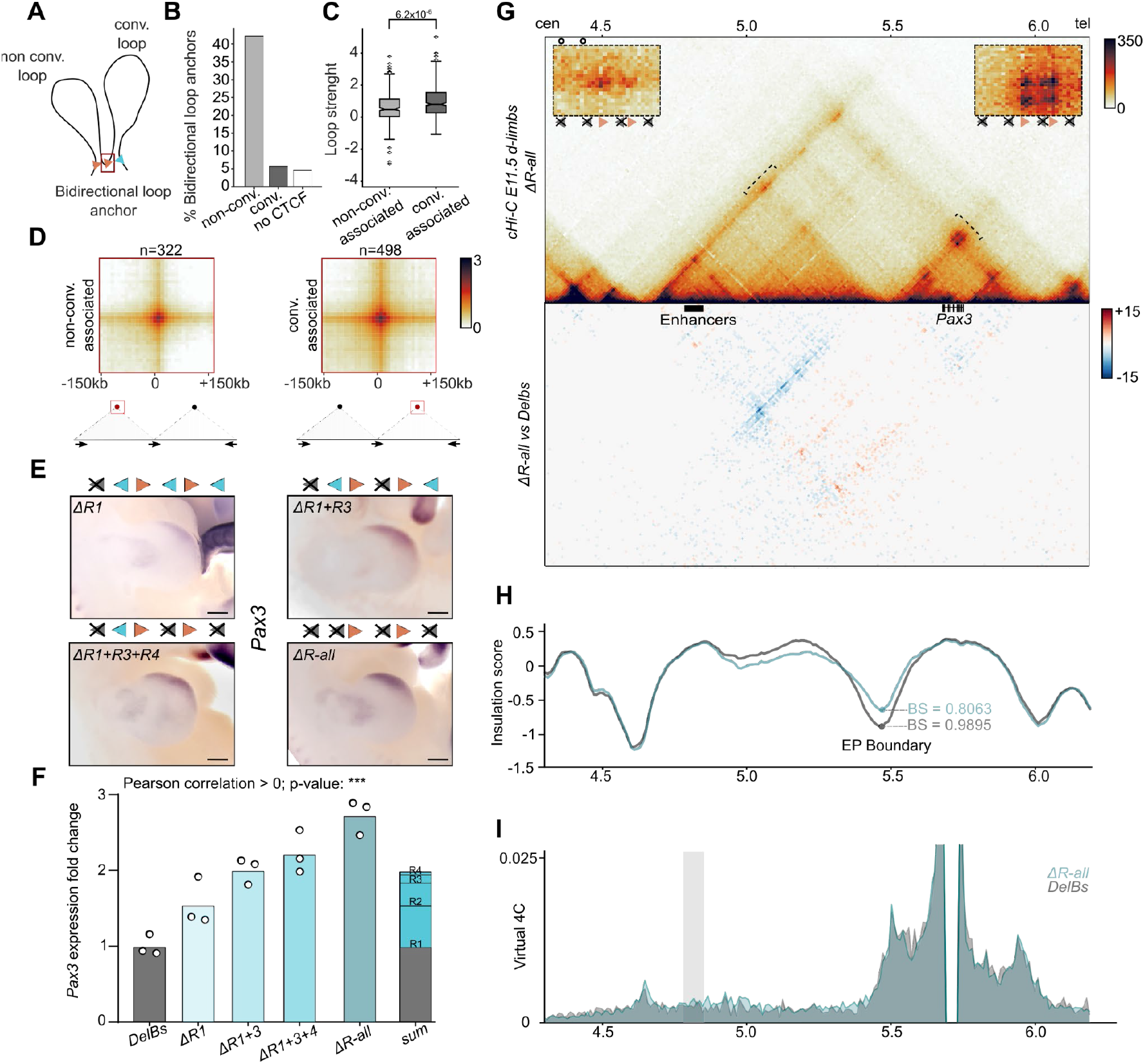
Formation of chromatin loops by non-convergently oriented CTCF binding sites. **A.** Schematic showing a hypothetical convergent loop (right) that indirectly generates a non-convergent loop in the opposite direction (left). **B.** Percentage of loop anchors that establish loops in both directions (dataset of mouse ES-cells from Bonev et al., 2017). Anchor categories: *convergent-only* (if they always contain CBS oriented in the same direction as their anchored loops), *non-convergent* (if they anchor at least one loop in a direction for which they lack a directional CBS) and *no-CTCF* (they do not contain any CBS). **C**. Loop strengths (calculated as in Flyamer et al. 2017, dataset of mouse ES-cells from Bonev et al., 2017) in pairs of convergent/non-convergent loops. The loops are classified in *non-conv. associated* (non-convergent loop that share the non-convergent anchor with a convergent loop established in the opposite direction-loop to the left in **A**-) and *conv-associated* (convergent loop that share one of the anchors with a non-convergent loop in the opposite direction -loop to the right in **A**-). **D**. Aggregated loop signal for categories described in **C**. Arrows in schematic represent CBS orientation. **E.** WISH shows *Pax3* expression in E11.5 forelimbs from CBS mutants. Arrowheads represent reverse (light blue) and forward (orange) oriented CBS. Crosses indicate deleted CBS. Note positive correlation between expanded *Pax3* misexpression and increase number of deleted CBS. Scale bar: 250μm. **F.** *Pax3* qPCR analysis in E11.5 limb buds from CBS mutants. Bars represent the mean and white dots represent individual replicates. Values normalized against *DelBs* mutant (ΔΔCt). Note positive correlation of *Pax3* misexpression with the increase in deleted CBS (Pearson correlation significantly >0; *** p-value ≤0,001). **G.** c-HiC maps from E11.5 mutant distal limbs at 10 kb resolution (top). Data mapped on custom genome containing the *DelBs* deletion. Insets represent a magnification (5kb resolution) of the centromeric (left) and telomeric (right) loops highlighted by brackets on the map. Gained or lost chromatin loops represented by full or empty dots, respectively. Subtraction maps (bottom) showing gain (red) or loss (blue) of interactions in mutants compared to *DelBs*. **H.** Insulation score values. Lines represent mutants. Dots represent the local minima of the insulation score at EP boundary for each mutant, also measured as boundary score (BS). **I.** Virtual 4C profiles with *Pax3* promoter as a viewpoint for the genomic region displayed in panel C. Light gray rectangle highlights *Epha4* enhancer region. Note increased interactions between *Pax3* promoter and *Epha4* enhancer in *ΔR-all* (blue) compared to *DelBs* mutants (gray).

To validate these findings *in vivo*, we sequentially retargeted our *ΔR1* mESCs to create a mutant that only retains the forward F1 and F2 sites. These CBS display a strong functionality according to their individual and combined deletions (**Fig. 2A and 2B**). Of note, both CBS are arranged in the same orientation, therefore the resulting mutant would lack the typical divergent CBS signature at the EP boundary. While generating this mutant, we also obtained intermediate mutants with a double deletion of the R1 and R3 sites (*ΔR1+R3*) and a triple deletion of R1, R3 and R4 (*ΔR1+R3+R4*), as well as the intended quadruple knockout lacking all reverse CBS (*ΔR-all*). WISH experiments in mutant limbs revealed an expanded *Pax3* expression pattern towards the posterior region of the limb, an effect that is more pronounced as the number of deleted CBS increases (**Fig. 3E**). Expression analyses by qPCR confirmed a significant increasing trend in *Pax3* misexpression levels across mutants (**Fig. 3F**; Pearson correlation > 0 p-value ≤ 2×10^-7^). These results, in combination with our observations on previous mutants, demonstrate that R2, R3 and R4 are indeed functionally redundant sites, despite the absence of measurable effects upon individual deletions (**Fig. 1B**). However, despite the increased expression, we noted that *Pax3* levels were only moderately increased (3-fold) compared to the increase in expression in mutants retaining reverse CBS (9-fold, *ΔF-all*). This is in stark contrast with the fact that *ΔR-all* mutants only retain two intact CBS in the forward orientation, while up to four CBS are still present in *ΔF-all* mutants. This suggests that the two forward CBS (F1 and F2) grant most of the insulator activity of the EP boundary. Thereby, these experiments confirm that the distinct functional characteristics of specific CBS can outweigh other predictive parameters of boundary function like the total number of sites.

As expected, the examination of cHi-C maps from *ΔR-all* mutant limbs revealed a clear partition between the *Epha4* and *Pax3* TADs (**Fig. 3G**), consistent with the reduced levels of *Pax3* misexpression. Boundary scores at the EP boundary were also only moderately reduced in comparison with the *ΔF-all* mutant, with only a 19% decrease compared to *DelBs* controls (**Fig. 3H,** compared to *ΔF-all* in **Fig.2D**). Accordingly, intra-TAD interactions modestly decreased while inter-TAD interactions increased, as observed for the interaction frequencies between the *Pax3* promoter and *Epha4* limb enhancers (**Fig. 3I**) Despite the multiple deletions, the telomeric chromatin loops remain unaffected and anchored by the F1 and F2 sites (**Fig. 3G; Supp. Fig. 8**). However, we noticed the persistence of centromeric chromatin loops in the complete absence of CBS with reverse orientation. Further examinations revealed that these loops become anchored by the F1 and F2 sites, despite their non-convergent forward orientation. In fact, we observe a higher contact intensity at F1, which would be the first CBS encountered by cohesin complexes sliding from the centromeric side. Therefore, our results in transgenic mice confirm our findings at the genome-wide level (**Fig. 3A, B and C**), demonstrating that CBS with robust insulator function can create chromatin loops independently of their CTCF motif orientation, most likely through a process of loop interference.

### The clustering of divergently-oriented CTCF binding sites is not a strict requirement for robust boundary insulation

Previous studies described the presence of divergent CBS clusters as a recurrent signature of TAD boundaries, suggesting a potential role on insulation (Gómez-Marín et al., 2015; Kentepozidou et al., 2020). While our analysis on mutants with non-divergent and only reverse CBS orientation (*ΔF-all*) showed a severe impairment of boundary function (**Fig 2C**), this was not the case for *ΔR-all* mutants, which only retain CBS in a forward orientation (**Fig 3F**). Indeed, the degree of *Pax3* misexpression in these mutants evidenced that insulation is more preserved than in *ΔR1+F2* mutants, which still conserve a divergent CBS signature (**Fig. 2C**).

This prompted us to explore the relation between the composition of CBS at boundaries and insulation strength. We examined available Hi-C datasets, classifying boundary regions according to different parameters of CBS composition (i.e., number and orientation) and calculating their boundary scores (**Fig. 4A**). Our analysis revealed that, for the same number of CBS, boundaries with a divergent signature generally display more insulation than their non-divergent counterparts. However, we observe that up to 6% of non-divergent boundaries display scores above 1.0, a value that we find to provide robust functional insulation to the EP boundary region (*DelBs* mutants; **Fig. 1C**). Manual inspection of specific loci showed that non-divergent boundaries with strong boundary scores present an evident TAD partition between adjacent domains in Hi-C maps. At these loci, insulation is not associated with prominent transcription or RNAPII occupancy at the boundary region. Furthermore, genes located at either side are unlikely to be coregulated (**Supp. Fig. 11**). These results, combined with our observation on CBS mutants, suggest that a divergent signature is not strictly required to form strong functional boundaries.

**Figure 4.**
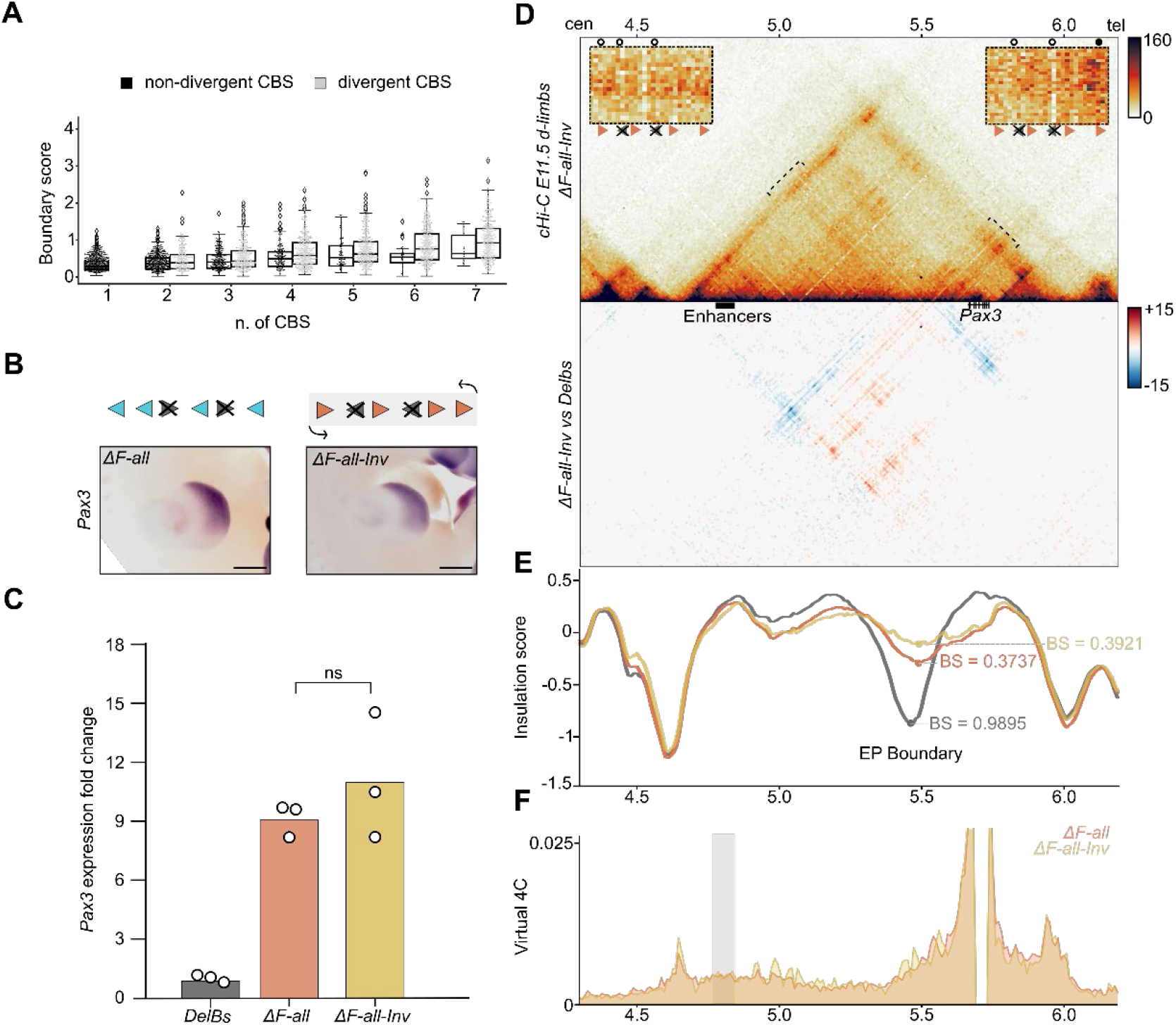
Non-divergent boundary signatures and effects of surrounding genomic context. **A.** Relation between Boundary Scores (BS) and the number of CBS for divergent and non-divergent boundaries in mouse ES-cell high-resolution Hi-C dataset (Bonev et al., 2017). The boxes in the boxplots indicate the median and the first and third quartiles (Q1 and Q3). Whiskers extend 1.5 times the interquartile range below and above Q1 and QR respectively. **B.** WISH shows *Pax3* expression in E11.5 forelimbs from CBS mutants. Arrowheads represent reverse (light blue) and forward (orange) oriented CBS. Crosses indicate deleted CBS. Light gray rectangle marks inverted region. Note similar *Pax3* misexpression pattern between *ΔF-all-Inv* and *ΔF-all* mutants. Scale bar: 500μm. **C.** *Pax3* qPCR analysis in E11.5 limb buds from CBS mutants. Bars represent the mean and white dots represent individual replicates. Values normalized against *DelBs* mutant (ΔΔCt). Note positive correlation of *Pax3* misexpression with the increase in deleted CBS (ns: non-significant). **D.** c-HiC maps from E11.5 mutant distal limbs at 10 kb resolution (top). Data mapped on custom genome containing the *DelBs* deletion and the inverted EP boundary. Insets represent a magnification (5kb resolution) of the centromeric (left) and telomeric (right) loops highlighted by brackets on the map. Gained or lost chromatin loops represented by full or empty dots, respectively. Subtraction maps (bottom) showing gain (red) or loss (blue) of interactions in mutants compared to *DelBs*. **E.** Insulation score values. Lines represent mutants. Dots represent the local minima of the insulation score at EP boundary for each mutant, also measured as boundary score (BS). **F.** Virtual 4C profiles with *Pax3* promoter as a viewpoint for the genomic region displayed in panel **C**. Light gray rectangle highlights *Epha4* enhancer region. Note similar interaction profile between *ΔF-all-Inv* (yellow) and *ΔF-all* mutants (orange).

### The orientation of large boundary regions has a limited impact on boundary insulation

Our analyses on individual mutants suggest a strong influence of CBS characteristics on boundary function. This effect can explain the prominent differences in insulation observed between mutants with only-reverse (*ΔF-all*) or only-forward (*ΔR-all*) sites. However, it is also plausible that the different genomic contexts at both sides of the EP boundary region could account for these functional differences. To evaluate this hypothesis, we generated a mutant that carries an inversion of the entire boundary region, on the *ΔF-all* background (*ΔF-all-Inv*) (**Fig. 4B**). Following the retargeting of our *ΔF-all* mESC, we confirmed the presence of two inverted alleles by qPCR (**Supp. Fig. 12**). Then, mutant embryos were generated from the modified mESC via tetraploid aggregation methods and subsequently analyzed.

WISH and qPCR experiments show that the pattern of *Pax3* expression is almost indistinguishable from the *ΔF-all* mutants, both spatially and at the quantitative level (**Fig. 4B and 4C**). The examination of cHi-C maps from *ΔF-all-Inv* mutants also revealed a partial fusion of the *Epha4* and *Pax3* TADs, similar to the spatial configuration of *ΔF-all* mutants (**Fig. 4D**). However, subtraction maps revealed qualitative differences related to the chromatin loops anchored at the EP boundary. Specifically, the inversion of the entire boundary region causes a redirection of chromatin loops, which now interact mainly with the telomeric *Pax3* boundary instead of the centromeric *Epha4* boundary. Nevertheless, these ectopic loops are mainly anchored by the R1 site, which preserves its marked functionality among the cluster. However, despite the observed local differences, the boundary scores indicate that boundary function is comparable between *ΔF-all-Inv* and *ΔF-all* mutants (**Fig. 4E**). This is also evident on virtual 4C profiles, which show a similar degree of interactions between *Pax3* and the *Epha4* enhancers (**Fig. 4F**). These results suggest that the orientation of entire boundary regions, as well as the differences in the surrounding genomic context, play a minor role in insulator function.

### Genomic distances can influence gene expression levels

To determine to what extent CTCF binding contributes to the function of the EP boundary region, we retargeted our mESCs to generate a sextuple knockout mutant where all CBSs are deleted (*ΔALL*). WISH experiments on tetraploid-derived embryos revealed a further expansion on the spatial pattern of *Pax3* misexpression, which covered the entire distal limb from the anterior to the posterior region. This expanded expression mirrors the pattern observed in *DelB* mutants, where the entire boundary region is deleted (**Figure 5A**). Expression analyses revealed that *Pax3* misexpression levels in *ΔALL* mutants exceed the combined sum of expression of the *ΔR-all* and *ΔF-all* mutants (**Fig. 5B**), again indicating the cooperative and redundant action of CBS in preserving insulation. Intriguingly, *ΔALL* mutants only reach 65% of the *Pax3* misexpression levels observed in the *DelB* mutants (**Figure 5B**), an effect that could be attributed to the 150Kb inter-CTCF region that differentiates both mutants.

**Figure 5.**
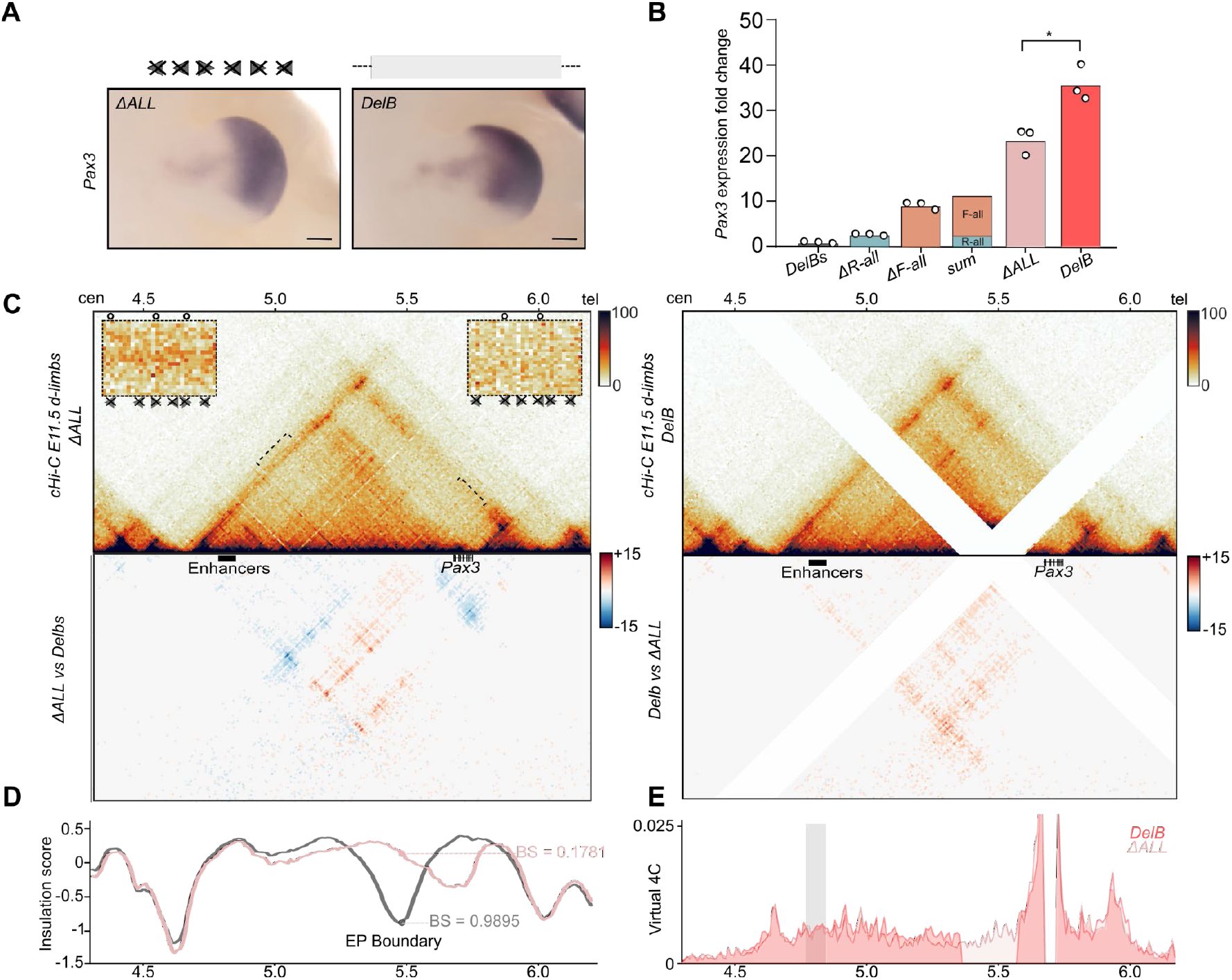
Contribution of CTCF binding to the insulation function of the EP boundary. **A.** WISH shows *Pax3* expression in E11.5 forelimbs from CBS mutants. Arrowheads represent reverse (light blue) and forward (orange) oriented CBS. Crosses indicate deleted CBS and gray rectangle represents deleted region. Note the similarities in expression pattern between mutants. Scale bar: 250μm **B.** *Pax3* qPCR analysis in E11.5 limb buds from CBS mutants. Bars represent the mean and white dots represent individual replicates. Values normalized against *DelBs* mutants (ΔΔCt). Note positive correlation of *Pax3* misexpression with the increase in deleted CBS (* p-value ≤ 0.05) **C.** c-HiC maps from E11.5 mutant distal limbs at 10 kb resolution (top). Data mapped on custom genome containing the *DelBs* deletion and the inverted EP boundary. Insets represent a magnification (5kb resolution) of the centromeric (left) and telomeric (right) loops highlighted by brackets on the map. Gained or lost chromatin loops represented by full or empty dots, respectively. Subtraction maps (bottom) showing gain (red) or loss (blue) of interactions in mutants compared to *DelBs*. **D.** Insulation score values. Lines represent mutants. Dots represent the local minima of the insulation score at EP boundary for each mutant, also measured as boundary score (BS). **E.** Virtual 4C profiles with *Pax3* promoter as a viewpoint for the genomic region displayed in panel **C**. Light gray rectangle highlights *Epha4* enhancer region.

To investigate the molecular cause of the reduced *Pax3* misexpression in *ΔALL* mutants, we performed cHi-C experiments in distal developing limbs (**Fig. 5C**). These experiments revealed a prominent fusion of the *Epha4* and *Pax3* TADs. This results from a severe disruption of the EP boundary, which displays a lower boundary score compared to the *DelBs* (**Fig. 5D;** 81% boundary score reduction) and a complete absence of anchored chromatin loops within the region. In fact, the interaction profile at the EP boundary is not different from other internal locations of the *Epha4* TAD (**Fig. 5C**). However, virtual 4C profiles from *ΔALL* mutants showed decreased interactions between *Pax3* and the *Epha4* enhancers, in comparison to *DelB* mutants (**Fig. 5E**) thus in agreement with the differences of *Pax3* misexpression observed between these mutants. ChIP-seq datasets for epigenetic marks did not reveal additional regions with regulatory potential within the 150 Kb region (**Supp. Fig. 3**), making it unlikely that the higher levels of *Pax3* misexpression in *DelB* mutants are caused by a deletion of regulatory elements. Taken together, these results suggest that enhancer-promoter distances might influence gene expression levels, thus providing a potential explanation for the reduced *Pax3* expression levels in *ΔALL* compared to *DelB* mutant embryos.

### Boundary insulation as a modulator of developmental gene expression and phenotypes

We previously reported that the misexpression of *PAX3* during early limb development can lead to a severe shortening of index and thumb fingers (brachydactyly), as observed in human patients carrying large deletions at the *EPHA4* locus and in their corresponding mouse models (*DelB*) (Lupiáñez et al., 2015). Therefore, our collection of mouse mutants provides a unique opportunity to study how boundary insulation strengths directly translate into developmental phenotypes.

To evaluate this aspect, we performed tetraploid aggregation experiments with several of our mutant mESC lines and obtained mutant fetuses at E17.5, a developmental stage where the limb defects are already observable (Lupiáñez et al., 2015). We performed alcian blue/alizarin red skeletal staining in mutant limbs and measured relative digit length as a proxy for the phenotype (**Fig. 6A and B**). First, we analyzed the *ΔR1* mutants, which display a moderate *Pax3* misexpression in the anterior region of the distal limb (**Fig. 1F**). A quantification of finger length ratios revealed that mutant limbs are indistinguishable from their corresponding controls. These results demonstrate that the detrimental effects of *Pax3* misexpression can be partially buffered, resulting in the development of normal limbs.

**Figure 6.**
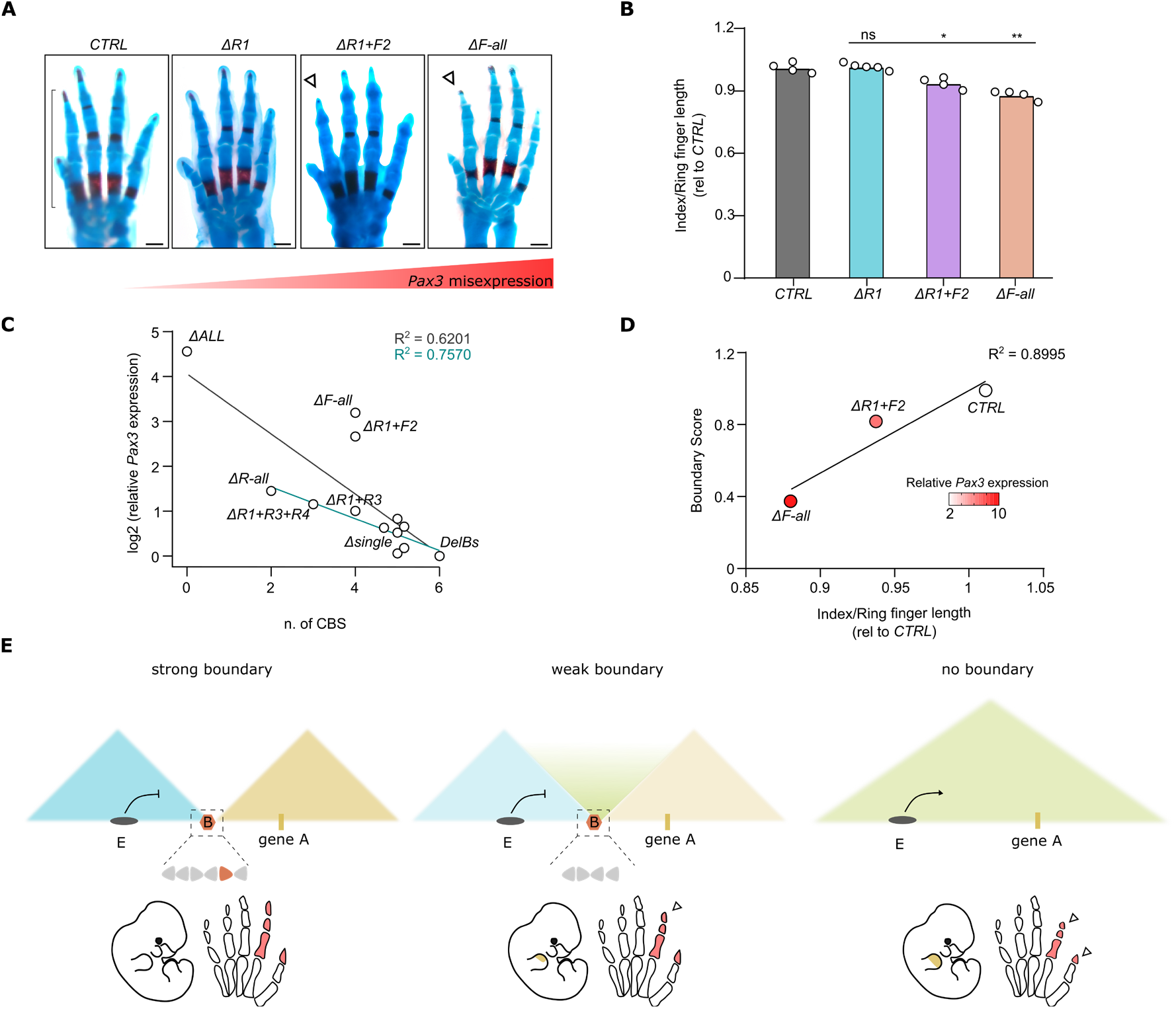
Boundary strength modulates developmental phenotypes. **A**. Skeletal staining of forelimbs from E17.5 mutant and control fetuses. White arrowheads indicate reduced index finger lengths. Black bracket shows the region of the finger measured for the quantification. Finger length correlates negatively with increased *Pax3* misexpression. Scale bar, 500μm. **B**. Index lengths relative to ring finger lengths in E17.5 mouse forelimbs. Bars represent the mean and white dots represent individual replicates. Values normalized on control animals (CTRL). (* p-value ≤ 0.05; ** p-value ≤ 0.01; ns: non-significant). **C.** Correlation between the number of remaining CBS at the EP boundary and the levels of *Pax3* expression in the different mutants described in this study. Pearson regression lines are shown together with R^2^ values, both for the whole collection of mutants (black) and discarding combined CBS deletions involving CBS with forward orientation (turquoise). **D.** Correlation and R^2^ between Boundary Scores and the brachydactyly phenotype penetrance measured as the Index to Ring finger length ratio for controls, *ΔR1+F2* and *ΔF-all* mutants. The color of the dots represents the level of *Pax3* misexpression in the limb as measured by qPCR. **E.** Model for boundary insulation as a quantitative modulator of gene expression and developmental phenotypes. Left, a strong boundary (B) efficiently insulates gene A from the enhancers located in the adjacent TAD (E). The boundary shows a cluster of CBS with different orientations represented with arrowheads. The colored arrow represents a CBS with prominent contribution to boundary function. Middle, the absence of specific CBS results in a weakened boundary, moderate gene misexpression (limb, indicated in yellow) and mild phenotypes (reduced digits, indicated in red and pointed by white arrowhead). Right, the absence of the boundary causes a fusion of TADs, strong gene misexpression and strong phenotypes.

Next, we analyzed the phenotypic effects of the *ΔR1+F2* mutant. In contrast to individual deletions, the combined deletion of R1 and F2 led to a moderate reduction of index digit length (**Fig 6A and B;** 6.3% compared to controls), consistent with the increased *Pax3* misexpression (**Fig. 2B**). This phenotype demonstrates that weakened boundaries can be permissive to functional interactions between adjacent TADs, resulting in developmental gene expression patterns and associated phenotypes. It is worth noting that the brachydactyly phenotypes of *ΔR1+F2* mutants occur despite an observable partition between the *Epha4* and *Pax3* TADs and across a boundary region that has relatively high boundary scores (**Fig. 2C and 2D**; boundary score=0.8). Further analyses on the distribution of boundary scores from ultra-high resolution Hi-C datasets (Bonev et al., 2017) revealed that many boundary scores fall within the range observed in our mutant collection (**Supp. Fig. 13**). Particularly interesting are the 40% of boundaries that display scores lower than 0.8, which according to our functional observations could be permeable for functional interactions across domains, according to our observations at the EP boundary. Nevertheless, insulation is not the only aspect that should be taken into account when evaluating enhancer-promoter relationships. Locus-specific features, such as the variability in enhancer strength or enhancer-promoter compatibility, might compensate for lower insulation levels.

Finally, we analyzed the *ΔF-all* mutants, in which the *Epha4* and *Pax3* TADs appear largely fused in cHi-C maps (**Fig. 2C**). In this case, the more severe disruption of TAD organization led to a prominent reduction of digit length (**Fig 6A and B;** 12% compared to controls), again consistent with the higher degree of *Pax3* misexpression observed in these mutants (**Fig. 2B**). Overall, these results illustrate how boundary insulation strength can serve as a modulator of gene expression and developmental phenotypes, by allowing permissive functional interactions between neighboring TADs.

## DISCUSSION

By combining genome-wide analyses and a comprehensive set of mouse mutants, we reveal principles of boundary function in *vivo*. Using the EP boundary region as a prototypical example, we demonstrate that CBS act redundantly and also cooperate to establish precise levels of regulatory insulation. On the one hand, the EP boundary function was increasingly compromised with the number of CBS mutations and remained almost unaffected upon individual deletions. On the other hand, combined mutants carrying either the F1 or F2 CBS deletions resulted in enhanced *Pax3* misexpression, thus escaping the general additive trend observed for the consecutive mutations of reverse oriented CBS (**Fig. 6C**). Therefore, boundary function appears to be highly determined by the characteristics of specific CBS, a parameter that can contribute more to effective domain insulation than the total number of sites or their orientation. We also observe that the stark differences in functionality between CBS often correlate with differences in CTCF occupancy, despite prominent exceptions like for R3. The latter suggests that additional factors may modulate CBS function. A recent study based on an *in vitro* insulator reporter assay revealed that flanking genomic regions can also contribute to CBS function, potentially serving as binding platforms for such additional modulators that are yet to be identified (Huang et al., 2020). Interestingly, this study also demonstrates that tandem CBS can display synergistic effects on insulation. However, we observe that CBS can compensate for the absence of each other to some extent, as combined deletions are required to compromise EP boundary insulation severely. Therefore, our results show that synergistic effects are negligible when the number of clustered CBS increases.

The combination of both cooperative and redundant functions have been also extensively described for other types of non-coding elements. For example, enhancer redundancy is a landmark of developmental gene regulation that confers phenotypic robustness and buffers against the detrimental effects of genetic perturbations (Frankel et al., 2010; Osterwalder et al., 2018). Several examples illustrate that enhancer elements can build cooperative regulatory networks that display striking differences on the activity of individual components, as we also report for CBS (Hay et al., 2016; Shin et al., 2016; Will et al., 2017). Therefore, our results suggest that boundary elements operate under biological principles that are not drastically different from other classes of genomic non-coding elements. This is particularly noticeable for CBS that do not show any apparent function upon individual deletion, but that can compensate for the loss of neighboring CBS. The redundancy of CBS seems to converge with additional buffering mechanisms that operate during development. For example, we observed that the moderate misexpression of *Pax3* resulting from the partial disruption of the EP boundary is not sufficient to cause abnormal phenotypes, demonstrating that other regulatory layers can compensate for detrimental gene effects. Therefore, developmental phenotypes are likely controlled by complementary “fail-safe” mechanisms that operate at multiple levels: the redundant action of different classes of non-coding elements, like enhancers and insulators, combined with downstream mechanisms that buffer fluctuations in gene expression.

The clustering of redundant CBS is a molecular signature that has been conserved across vertebrate evolution (Gómez-Marín et al., 2015; Vietri Rudan et al., 2015). This signature might have contributed to the conservation of relevant TAD boundaries across species, despite a constant turnover of individual CBS (Kentepozidou et al., 2020). Previous studies suggest that the evolutionary loss of individual CBS at clustered regions might have a low impact on the expression of nearby genes (Kentepozidou et al., 2020). However, the striking differences in functionality between individual sites might also result in differential selective pressures over CBS, thus causing evolutionary remodeling to occur preferentially at sites with reduced function and leading to minor changes in boundary insulation. Nevertheless, we also demonstrate that individual CBS deletions can “permeabilize” boundary regions and result in slight variations on gene expression that might be difficult to identify in quantitative analyses. Thus, it is conceivable that boundary regions, despite a robust evolutionary conservation, might also serve as a substrate for gradual changes in gene expression and corresponding phenotypes.

The transcriptional changes upon boundary perturbation show a marked tissue and temporal specificity. This suggests that selective constraints on insulation are mostly determined by the specific activity of regulatory elements, as well by the responsiveness of genes located at the opposite side of a boundary. Thus, it is expected that a global depletion of 3D chromatin organization in individual cell types may only results in moderate changes in transcription, due to a reduced number of active enhancers, as reported in *in vitro* studies (Nora et al., 2017; Rao et al., 2017). Yet, even a limited number of dysregulated genes can be sufficient to induce severe developmental defects *in vivo* (Franke et al., 2016; Lupiáñez et al., 2015). Further studies that abolish 3D chromatin organization across multiple tissues and at developmental stages will be essential to ultimately determine the *in vivo* relevance of this process.

Our study also offers novel mechanistic clues concerning TAD boundary formation. While divergent CBS signatures are associated with stronger boundaries on average, they are not a strict requirement for robust boundary function. Non-divergent boundaries can also display boundary scores that go above those values reported to be functionally robust for the EP boundary. While the underrepresentation of non-divergent boundaries across the genome could be interpreted as a sign of decreased functionality, we cannot ignore the fact that for an increasing number of CBS the probabilities of not having a divergent signature increase exponentially. Additionally, we observe that many CBS boundaries with non-divergent signatures that are formed by non-convergent loops paired to convergent ones. Such configuration can be mechanistically explained by a process of loop interference, where the persistent anchoring of a cohesin complex might stall additional complexes. Therefore, our results constitute an experimental validation for a scenario that is predicted by the loop extrusion model (Fudenberg et al., 2016; Sanborn et al., 2015) and that might be linked to the existence of robust chromatin boundaries with non-divergent CBS signatures.

While it is well proven that boundaries can effectively constrain interactions between neighboring TADs, whether insulation can be considered an absolute property from a functional and regulatory perspective remained unclear (Chang et al., 2020). Single-cell (Flyamer et al., 2017; Nagano et al., 2017) and super resolution microscopy approaches (Bintu et al., 2018; Szabo et al., 2018) demonstrated that chromatin interactions in individual cells are stochastic and, in some instances, ignore the presence of boundaries that are well-defined in bulk data. In light of these studies, our results reinforce the premise that boundary insulation should be considered as a quantitative property, as enhancer-promoter cross-talk and gene activation are largely proportional to boundary insulation strength (**Fig. 6D**). Further, boundary insulation appears to be also influenced by genomic distances, as illustrated in our comparison between *DelB* and *ΔALL* mutants where the presence of a 150 Kb region with no observable chromatin loops causes a 30% reduction in *Pax3* activation. A complementary observation was also described for inversions at the *Shh* locus, where reducing the distances between the ZRS enhancer and the *Shh* gene was sufficient to overcome boundary insulation and cause ectopic gene activation (Symmons et al., 2016). In any case, insulation strength emerges as a key feature of boundary function, which can effectively modulate gene activation and phenotypes (**Fig. 6E**). In turn, these two parameters would induce further developmental and evolutionary constraints that would influence the strength of boundary insulation. Therefore, we uncover that chromatin boundaries are modular and multicomponent genomic regions subjected to several principles that govern their regulatory logic. These principles should be broadly applicable to other loci, thus facilitating their functional interpretation in different developmental contexts. Such knowledge might help to bridge the gap between the information encoded in the 3D structure of genomes and the biological processes subjected to their control.

## MATERIAL AND METHODS

### Generation of CBS mutant mice

Mutant mESCs were obtained following an already described method (Kraft et al., 2015). All CBS deletions were generated using only one single guide RNA (sgRNA) designed in close proximity to the binding motif using the website Benchling (https://www.benchling.com/), with the only exception of *ΔR2* and *ΔF_ALL_Inv* that were generated by using a pair of sgRNA. The guide sequences (listed in **Supp. Table 1**) were then cloned in px459 CRISPR/Cas9 vector (Addgene Cat. N. 62988), previously digested with Bbs1. *Delbs* mESCs (4×10^5^) were seeded on a layer of inactive CD1-feeders and cultured with standard ES culture conditions. Cells were transfected using FuGene HD reagents (Promega) and 8 μg of each px459 vector containing the sgRNA of interest, following the kit guidelines. Twenty-four hours post-transfection, cells were split on puromycin resistant DR4-feeders and selected with puromycin for 48 hours. After selection, mESC colonies were left to recover for 3-5 days, then individual clones were picked and transferred to 96-well plates coated with CD1-feeders. After a few days, when cells had reached confluency, they were split in three; two parts were frozen and one part was left growing more for DNA isolation without feeders. Every clone was genotyped by PCR (MangoTaq, Bioline, Cat.N. 25033) (primers listed in **Supp. Table 2**) and positive clones were selected only if they showed two distinguishable bands on agarose gel, representing two different deleted alleles. Homozygous deletions were further confirmed by Sanger sequencing. Positive clones were thawed, expanded and genotyped again to further confirm the genotyping results. In order to obtain the combined CBS deletions, some individual CBS mutants were re-targeted following the procedure described.

The engineered cells were successively used to generate embryos by tetraploid aggregation methods (Artus and Hadjantonakis, 2011; Kraft et al., 2015). Mice were handled according to institutional guidelines under an experimentation license (G0111/17) approved by the Landesamt fuer Gesundheit und Soziales (Berlin, Germany) and housed in standard cages in a specific pathogen-free facility.

### Whole-mount in situ hybridization (WISH)

E11.5 mouse embryos were dissected in 1X PBS and fixed overnight in 4% PFS/PBS at 4°C. The following day, embryos were washed twice (10 minutes each) in 1X PBS/DEPC water and gradually dehydrated with different Methanol dilutions in PBS/DEPC water (25%, 50%, 75%) at 4°C for 30 minutes. Finally, embryos were washed twice (10 minutes each) with 100% Methanol and stored at −20°C. *Pax3* digoxigenin (DIG) - labeled antisense riboprobes were transcribed from linearized gene-specific probes (PCR DIG probe Synthesis Kit, Roche). WISH experiment was performed as follows. Embryos were re-hydrated stepwise in 75%,50%,25% Methanol/PBS-Tween20 (PBST), washed twice in PBST (10 minutes each wash), bleached on ice in 6% hydrogen peroxide/PBST for 1 hour and washed again in PBST. Embryos were then digested using Proteinase K (Sigma-Aldrich, REF 03115836001) (10mg/mL) for 3 minutes. Proteinase K was stopped by washing the embryos twice with Glycine (2mg/mL) in PBST. Then, embryos were washed 5 times in PBST and finally re-fixed for 20 minutes in 4% PFA/PBS, 0.2% glutaraldehyde and 0.1% Tween at room temperature. After further washing steps with PBST, embryos were incubated at 68°C in L1 Buffer (50% deionized formamide, 5X SSC, 1% SDS,0.1% Tween20 in DEPC water; pH4.5) for 10 minutes. For pre-hybridization, embryos were incubated in H1 Buffer (L1 Buffer with 0.1% tRNA and 0.05% heparin) for 2 hours at 68°C. Before the hybridization, *Pax3* DIG-probes were diluted in H1 Buffer and denatured for 10 minutes at 80°C. Embryos were finally incubated overnight at 68°C in H2 Buffer (H1 Buffer with 0.1% tRNA, 0.05% heparin and 1:100 *Pαx3*-DIG probes). The day after, in order to remove the unbound probes embryos went through several washing steps with pre heated (68°C) L1, L2 (50% deionized formamide, 2X SSC pH4.5, 0.1% Tween20 in DEPC water, pH4.5) and L3 (2X SSC pH4.5, 0.1% Tween20 in DEPC water, pH4.5). 3 washes per buffer, 30 minutes each, were performed. Embryos were cooled down to room temperature and washed in 1:1 L3 Buffer/RNAse solution (0.1M NaCl, 0.01M Tris pH 7.5, 0.2% Tween20, 10mg/mL RNAse A in H2O) for 5 minutes. Afterwards, embryos were incubated twice for 30 minutes at 37°C with RNAse solution, followed by 5 minutes incubation at room temperature with 1:1 RNAse solution/TBST-1 (140mM NaCl, 2.7mM KCl, 25mM Tris-HCl, 1% Tween20, pH7.5). After 3 washes in TBST1 (each of 5 minutes), embryos were incubated in blocking solution (TBST1 with 2% calf-serum, 0.2% bovine serum albumin) for 2 hours shaking at room temperature. Embryos were incubated overnight with anti-DIG antibody conjugated to alkaline phosphatase (1:5000) (no. 11093274910, Roche) at 4°C. After the overnight incubation, the unbound antibody was washed out with 8 times at room temperature (30 minutes each) with TBST 2 (TBST with 0.1% Tween 20 and 0.05% levamisole–tetramisole) and finally left at 4°C overnight. On the last day, embryos were stained after 3 equilibration washes of 20 minutes in AP buffer (0.02 M NaCl, 0.05 M MgCl_2_, 0.1% Tween 20, 0.1 M Tris–HCl and 0.05% levamisole– tetramisole in H_2_O), followed by staining with BM Purple AP Substrate (Roche). Embryos were then washed twice in alkaline phosphatase buffer, fixed in 4 % PFA/ PBS/ 0,2 % glutaraldehyde and 5mM EDTA and stored at 4°C. The stained embryos were imaged using a Zeiss Discovery V12 microscope and Leica DFC420 digital camera.

### Quantitative polymerase chain reaction (qPCR)

E11.5 mutant forelimb buds were dissected in 1X PBS, collected and snap frozen in liquid nitrogen. Tissue was dissolved in RLT with the help of syringes and RNA extracted following the guidelines of RNeasy Mini Kit (Qiagen). Reverse transcription was performed using Applied Biosystems_High-Capacity cDNA Reverse Transcription Kit (Cat. No. 4368814) following the manufacturer instructions and using 500ng of RNA as input material. qPCR was then performed for at least 3 biological replicates using Biozym Blue S’Green qPCR Mix Separate ROX (No. 331416XL) on QuantStudio 7 Flex Real-Time PCR System from Applied Biosystems. *Pax3* fold change was calculated from ΔCt using *Gapdh* as housekeeping gene (2^-(ΔCt)). ΔΔCt was then calculated using *DelBs* mean as a reference value.

### ChIP-seq

MEFs depleted mutant mESCs (5×10^6^) were washed twice washed with 1X PBS, dissociated with 1mL Trypsin and centrifuged for 5 minutes at 1100 rpm at room temperature. Cell pellet was resuspended in 11.7 mL of 10% FCS and then fixed by adding 325 μL of 37% Formaldehyde (Sigma-Aldrich) (final 1% FA) and incubated for 10 minutes at room temperature, while rotating. To stop the fixation process, the reaction was quenched on ice by adding 1 mL of 1.425M Glycine. Nuclei extraction was performed by adding 5 mL of ice-cold lysis buffer (10mM Tris HCl pH 7.5, 10 mM NaCl, 5 mM MgCl2, 0.1 mM EGTA, 1X Protease Inhibitor (Roche Ref. 5892791001) in Milli-Q Water). Extracted nuclei were then collected by centrifugation at 460g for 5 minutes at 4°C, washed with 1X PBS, snap frozen and stored at −80 °C or further processed using iDeal ChIP for Transcriptional Factors Kit (Diagenode) (Cat. N. C01010055). Briefly, cell nuclei were resuspended in 300μl of Shearing Buffer and Chromatin was sheared using Diagenode Bioruptor in order to achieve a fragment size ranging from 200-500 bp. Immunoprecipitation was done using 15-20 μg of DNA and 1 μg of CTCF Ab (Diagenode:C15410210) and all steps were performed following the manufacturer instructions. ChIP-seq libraries were prepared using the NEBNext Ultra II Library Prep Kit for Illumina. Input material ranged from 500pg to 15ng of immunoprecipitated DNA and processed according to the kit guidelines (NEBNext End Prep, Adaptor Ligation, PCR enrichment of Adaptor-Ligated DNA using NEBNext Multiplex Oligos for Illumina). Clean up and size selection were performed with AMPure beads (NEB). Library was sequenced with 30 millions of single end read of 75 nt on HiSeq4000 or NovaSeq platform.

### Capture-HiC

E11.5 mouse distal limb buds from homozygous mutants were microdissected in 1X PBS, resuspended and incubated in 1ml pre-warmed Trypsin for 5-10 minutes at 37°C. Trypsin was blocked by adding 5mL of 10% FCS/PBS. The tissue was further dissociated to make a single-cell-suspension by using a 40μm cell-strainer (Product N. 352340) and finally centrifuged at 1100 rpm for 5 minutes at room temperature. The pellet was then resuspended in a 2% PFA (in 10% FCS/PBS) fixation solution and incubated at room temperature for 10 minutes while tumbling. To stop the fixation process, the reaction was quenched on ice by adding Glycine (final concentration 125 mM) and centrifuged at 400g for 8 minutes at 4°C. Nuclei extraction was performed by adding 1.5mL of ice-cold lysis buffer (50mM Tris HCl pH 7.5, 150 mM NaCl, 5 mM EDTA, 0.5% NP-40, 1.15% Triton X-100, 25X Protease Inhibitor in Milli-Q Water). Extracted nuclei were then collected by centrifugation at 750g for 5 minutes at 4°C, washed with 1X PBS, snap frozen and stored at −80 °C. The 3C library was achieved by a DpnII digestion, a re-ligation of the digested fragments, de-crosslinking and DNA purification and further processed using SureSelectXT Target Enrichment System for the Illumina Platform (Agilent Technology). 200 ng - 3 μg of input material was sheared using Covaris Sonicator and the following parameters: duty cycle: 10%, intensity: 5, cycle per burst:200, time: 6 minutes, temperature: 4°C. Sheared DNA was then processed following the kit guidelines (end repair, dA-tailing, adaptor ligation, PCR enrichment of Adaptor ligated DNA, DNA purification, hybridization and capture). The hybridization was performed using SureSelect XT Custom RNA probes library (Cat # 5190-4836) designed on the genomic region mm9 chr1:71,000,000-81,000,000. The capture was performed using Streptavidin-Coated Beads (Invitrogen). PCR enrichment and sample indexing were done following Agilent instructions. Capture libraries were sequenced with 400 millions of 75-100bp paired-end reads on HiSeq4000 or NovaSeq platforms

### Skeletal preparation

E17.5 mouse fetuses were dissected in 1X PBS, sacrificed and then kept for 1 hour in water and later incubated in 65°C hot water for 1 minute. Skin and organs were removed mechanically with the help of forceps. Prepared fetuses were further processed with different solutions and serial overnight incubations at room temperature. On day 1, fetuses were fixed overnight with 100% Ethanol, while oscillating. On day 2, in order to stain for cartilage, 100% ethanol was replaced by Alcian Blue solution and samples incubated ove-night (150 mg/L Alcian Blue 8GX (Sigma-Aldrich) in 100% Ethanol and Acetic Acid Glacial). In the following days, Alcian Blue solution was replaced first with Ethanol 100% (day 3), then with 0.2 % KOH for digesting the tissues (day 4), with Alizarin Red to stain membranous bones (50 mg/mL Alizarin Red in 0.2% KOH) (day 5) and finally with 0.2% KOH again to finalize tissue digestion (day6). On day 6, they were placed in 25% Glycerin in Milli-Q Water for imaging acquisition. Stained fetuses were imaged using Zeiss Discovery V.12 microscope and Leica DFC420 digital camera. Fetuses were long-term stored in 60% Glycerin solution.

### *Capture-HiC* analysis

Paired-end reads from all the Capture-C experiments were aligned using bwa mem local aligner (Li and Durbin, 2010) to a custom reference genome encompassing the captured-region (chr1:71-81Mb of the mm9 assembly) with the region corresponding to the baseline *DelBs* mutation deleted (chr1:76,388,978-77,858,974). There was one exception, the *ΔF-all-Inv* mutant Capture-C, in which a different version of the genome was used to account for the inverted coordinates (chr1:77,861,422-78,062,382). The rest of chromosomes, including the remaining chr1, were kept in the custom reference genome to be able to distinguish not-uniquely mapped reads. Then, following the 4DN consortium recommendations, the resulting bam files were parsed with the pairtools suite (https://github.com/mirnylab/pairtools) to produce 4DN format files containing pairwise interactions. Briefly, bam files were parsed using pairtools parse. Then, not-uniquely mapped reads were filtered out using pairtools select (selecting UU, UR and RU pairs). Subsequently, pairs of reads were sorted and duplicated pairs were removed using pairtools sort and pairtools dedup respectively. Finally, dangling-ends were filtered out using a custom python script available in the gitlab repository. Filtered 4DN formatted pairs of interactions were then used to construct Knight-Ruiz normalized Hi-C matrices in hic format with Juicer (Durand et al., 2016). Such hic files were further visualized and analyzed with FAN-C (Kruse et al., 2020) and custom python code also available in our gitlab repository. Briefly, insulation scores, boundaries and boundary scores were calculated as described elsewhere (Crane et al., 2015) using the dedicated FAN-C functions through the FAN-C Api. Subtraction matrices were calculated as described elsewhere (Bianco et al., 2018) with minor modifications. Briefly, first the coverage of the matrices to be subtracted was equalized dividing by the total number of reads. Then, the two matrices were subtracted element-wise and each value of the subtraction was converted to a z-score taking into account the rest of values belonging to the same sub-diagonal (corresponding to interactions happening at equivalent genomic distances). Virtual 4C tracks were visualized and quantified using custom python and R scripts (available). WT, *DelBs* and *DelB* cHi-C raw reads were downloaded from GEO (GSE92291) (Bianco et al., 2018)

### HiC analysis

#### Data retrieval

Already processed hic files (Durand et al., 2016) from high resolution Hi-C datasets in mouse stem cells, neural progenitors and cortical neurons (Bonev et al., 2017) were obtained from the Juicebox repository (see index in hicfiles.tc4ga.com/juicebox.properties). CTCF ChIP-seq datasets from matching cell types were downloaded from GEO (see GSE). Hi-C data from mouse embryonic proximal and distal forelimbs (Rodríguez-Carballo et al., 2017) were also downloaded from GEO (see GSE101715) in validPairs format and subsequently converted to hic files using Juicer. Matching CTCF ChIP-seq data were obtained from GSE101714.

#### Boundary analysis

Insulation scores, boundaries and boundary scores (Crane et al., 2015) were calculated with FAN-C (Kruse et al., 2020) using Knight-Ruiz (KR) normalized matrices at 25kb resolution with a window size parameter of 250kb. Boundaries located in the vicinity (+-125kb) of extremely low mappable regions were filtered out. Low mappability regions were defined using a gaussian mixture model on the marginal counts of the raw Hi-C matrices (further details and masked regions available in the gitlab repository). CBS were predicted using CTCF peaks from matching ChIP-seq datasets, and CBS orientation was inferred using FIMO (Grant et al. 2011, using the flags --bfile --motif-- --max-stored-scores 1000000 and the CTCF PWM from JASPAR, background estimated using MEME fasta-get-markov utility). The highest scoring motif from each peak was retained for further analysis. For the mouse ES-cells dataset, the total number of CBS and the total number of divergent CBS pairs was then calculated for each boundary including a 100kb long flanking region call using BEDTools (Quinlan and Hall, 2010).

#### Loop analysis

We calculated loops using CPU hiccups (Durand et al., 2016) with the flags (-m 512 -r 5000,10000,25000 -k KR -f .1,.1,.1 -p 4,2,1 -i 7,5,3 -t 0.02,1.5,1.75,2 -d 20000,20000,50000) in 5kb,10kb and 25kb KR-normalized matrices for the mouse ES-cells dataset (Bonev et al., 2017). Loop anchors were intersected with the CBS information obtained as described in the Boundary analysis section using BEDTools. Then, loop anchors were classified accordingly in convergent-only (loop anchors that display at least one CBS oriented in the direction of all the loops they are engaged), non-convergent (loop anchors that are engaged in at least one loop that is formed despite lacking any CBS oriented in that direction) and non-CTCF (loop anchors that do not display any CBS). CTCF loops were subsequently classified in two categories according to the nature of their anchors: convergent (loops formed by anchors displaying convergently oriented CBS) and non-convergent (if not). Convergent loops were further subdivided in single-sided convergent (if both anchors only engage in loops in the same direction), double-sided convergent (if at least one of the anchors engage in a convergent loop in the opposite direction) and convergent associated (if at least one of the anchors engage in a non-convergent loop in the opposite direction). Non-convergent loops were also subdivided in simply Non-convergent and non-convergent associated (if at least one of the anchors is engaged in a convergent loop in the opposite direction). Loop strengths were calculated for each set of loops as previously proposed (Flyamer et al., 2017) using the dedicated FAN-C function (Kruse et al., 2020). Hi-C signal aggregates over the different loop categories were also calculated using FAN-C and 10Kb matrices.

## ACKNOWLEDGMENTS

We thank the sequencing core, transgenic unit and animal facilities of the Max Planck Institute for Molecular Genetics and Max Delbrück Centre for Molecular Medicine for technical assistance. We thank Salaheddine Ali and Myriam Hochradel for their support in the preparation of capture Hi-C libraries. We thank Christina Paliou, Martin Franke, Marc A. Marti-Renom and members of the Lupiañez lab for their valuable input and comments on the manuscript.

## AUTHOR CONTRIBUTIONS

C.A., R.D.A. and D.G.L. conceived the study and designed the experiments. C.A. performed most experiments with support of A.B. and F.M.R. R.D.A. performed bioinformatics analysis. J.J., R.K. and L.W. performed tetraploid aggregation. G.C. and N.B. performed *in situ* hybridizations. C.A., R.D.A. and D.G.L. wrote the manuscript with input from all authors.

