## Supplementary Figures for "*In vivo* dissection of a clustered-CTCF domain boundary reveals developmental principles of regulatory insulation"

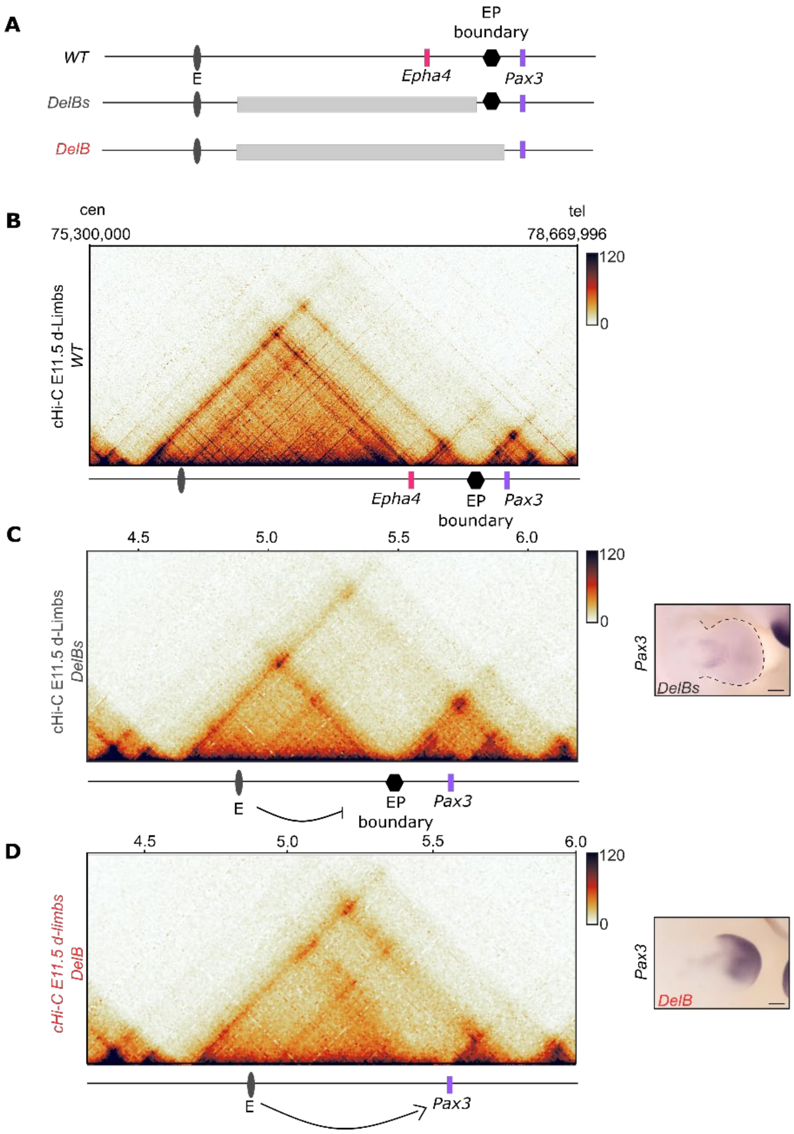


**Supp. Fig.1 Structural and molecular comparison between *Delbs* and *DelB* mutants.**

**A.** Schematic shows *Epha4* enhancers (in dark gray), *Epha4* gene (in pink), *Pax3* gene (in purple), EP boundary (in black) and genomic rearrangements in the *DelBs* and *Delb* mutants (light gray rectangles).

**B-D.** Capture Hi-C (cHi-C) maps in E11.5 limbs in WT (**B**) *DelBs* (**C**) and *DelB* (**D**) mutants (data from Bianco et al., 2018). Genomic coordinates in C and D correspond to custom *DelBs* and *DelB* genomes for the captured region, respectively. *Pax3* WISH (right panel). Note how the presence of the EP boundary is sufficient to block the functional interaction between the *Epha4* enhancers and the *Pax3* gene, thus preventing its misexpression. Cen, centromeric. Tel, telomeric. d-Limbs, distal limbs. Scale bars, 250 µm.

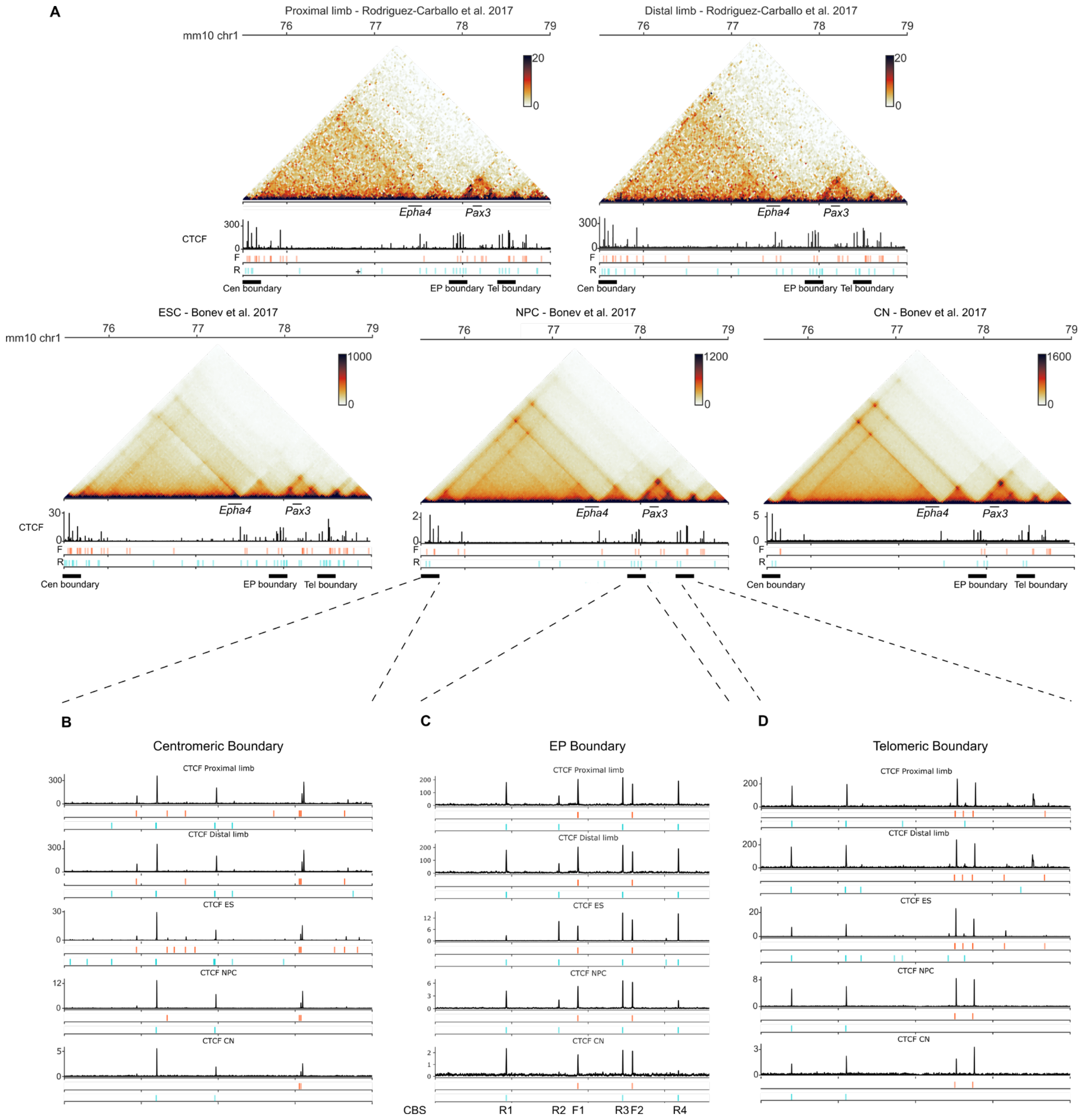


**Supp. Fig. 2. *Epha4-Pax3* (EP) boundary is a constitutive boundary with stable binding of divergently oriented CTCF.**

**A.** Hi-C maps at 25kb resolution and CTCF ChIP-seq tracks around the EP boundary locus from 5 different sources are shown. CTCF motifs inside CTCF peaks in a forward (F) or reverse (R) orientation are depicted below the ChIP-seq track in red and blue respectively. Motifs were calculated using FIMO (Grant et al., 2011, see methods). The first two datasets are proximal and distal embryonic forelimbs respectively (Rodriguez-Carballo et al., 2017). The other three datasets correspond to the high-resolution datasets in mESC, neural progenitor cells and cortical neurons from Bonev et al. 2017. **B-D.** Close-ups of the corresponding CTCF ChIP-seq experiments in the centromeric (Cen), *Epha4-Pax3* (EP) and telomeric (Tel) boundaries respectively.

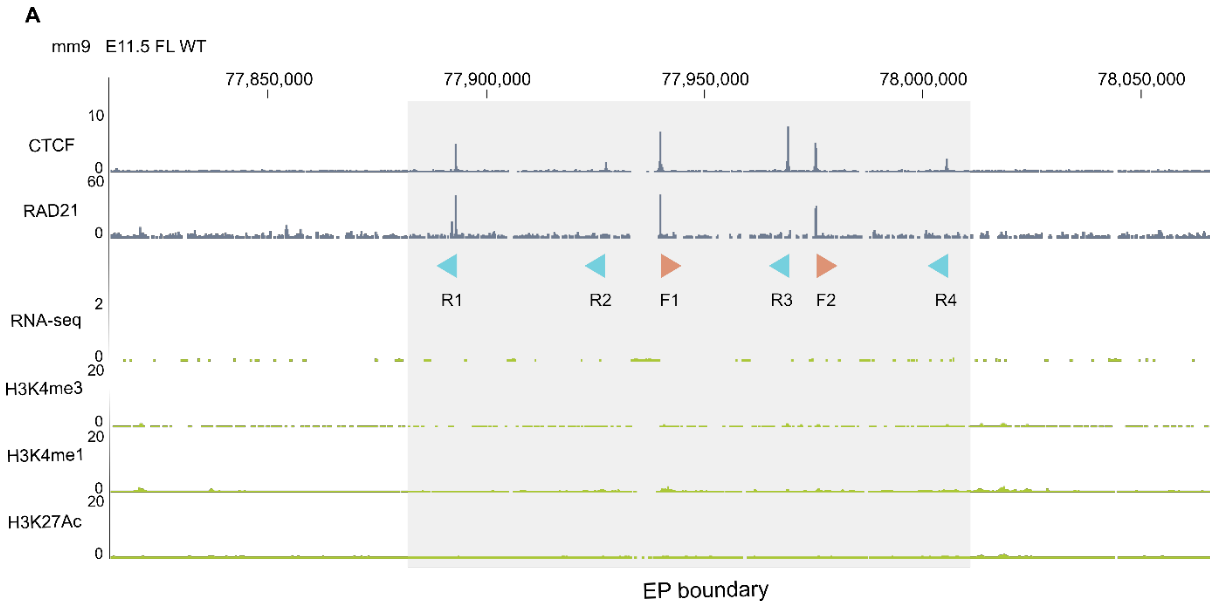


**Supp. Fig.3 Epigenetic landscape at the *Epha4-Pax3* locus.**

**A.** Genome browser tracks showing CTCF, RAD21, H3K4me3, H3K27Ac, H3K4me1 ChIP-seq and RNA-seq of E11.5 WT mouse forelimbs (FL) (data from Andrey et al., 2017). Note how the EP boundary is occupied by CTCF and RAD21, but shows no presence of the other histone marks nor of active transcription. EP boundary is indicated by the gray box. Light blue and orange arrowheads represent reverse (R) and forward (F) oriented CBS, respectively.

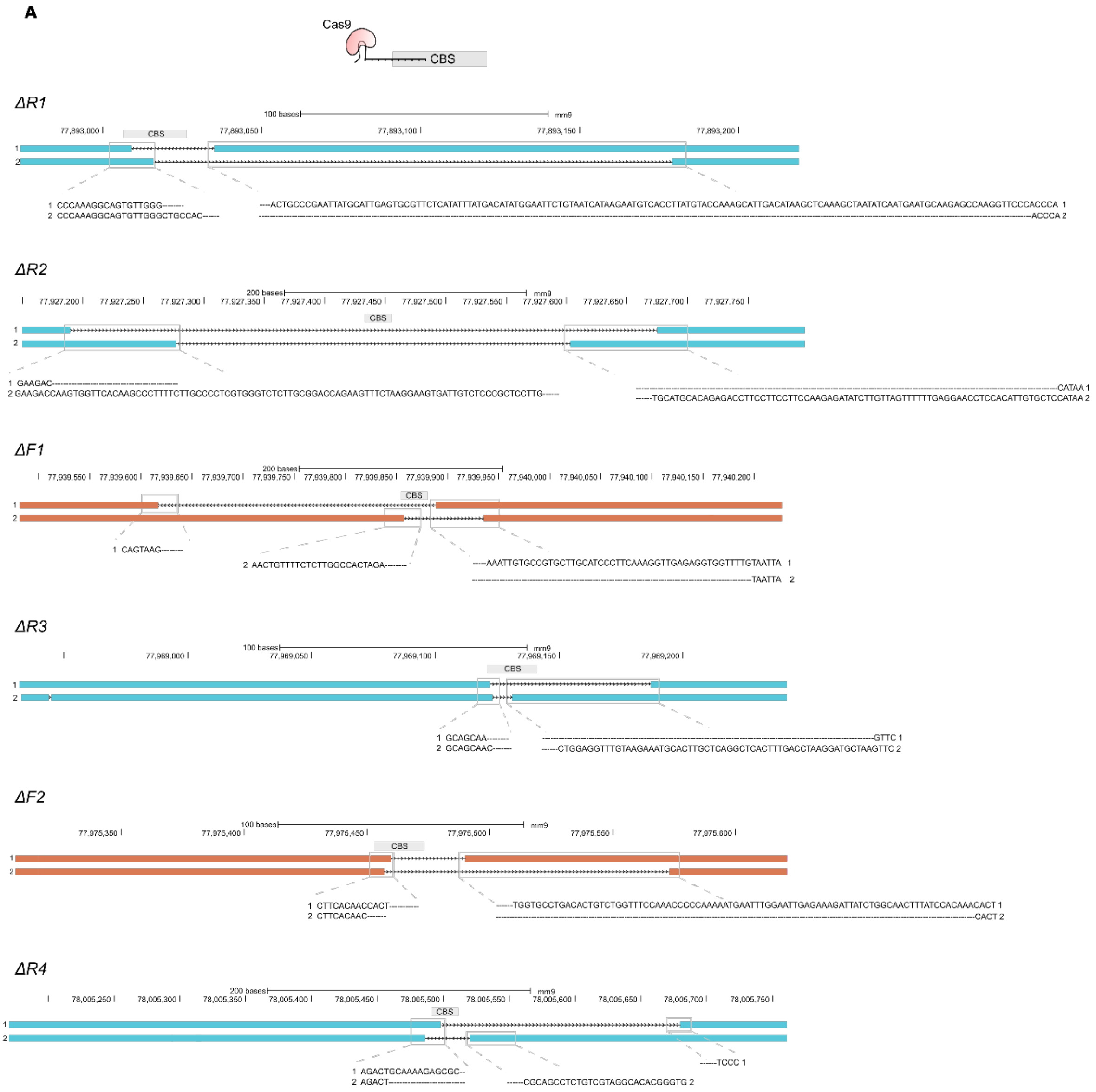


**Supp.Fig.4 CBS individual deletions generated by CRISPR/Cas9.**

**A.** Sanger sequencing reads schematics showing the deletion profiles for the six individual CBS mutants for the two alleles. CBS positions are specified. Deleted allelic regions are represented as black dashed lines. Intact allelic regions are indicated with light blue (reverse CBS) and orange (forward CBS) bars. Close-ups show the exact breakpoints sequences.

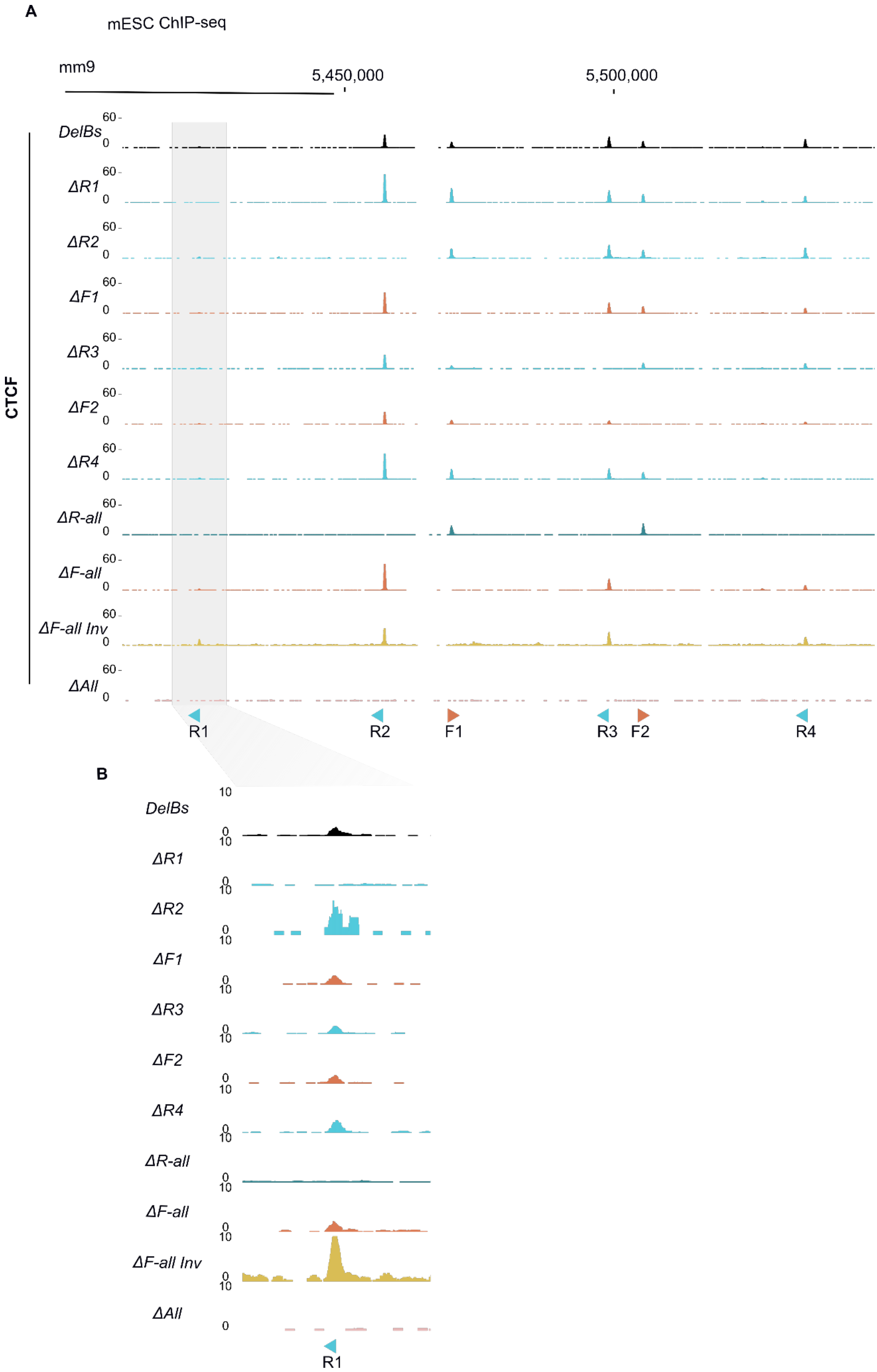
A
mESC ChIP-seq
mm9
5,450,000
5,500,000
CTCF
DelBs
ΔR1
ΔR2
ΔF1
ΔR3
ΔF2
ΔR4
ΔR-all
ΔF-all
ΔF-all Inv
ΔAll
60
0
R1
R2
F1
R3
F2
R4
B
10
0
DelBs
ΔR1
ΔR2
ΔF1
ΔR3
ΔF2
ΔR4
ΔR-all
ΔF-all
ΔF-all Inv
ΔAll
R1

**Supp. Fig. 5. Targeted CBS deletions completely abrogate CTCF binding in the intended loci.**

**A.** CTCF ChIP-seq experiments performed in the different mESC mutant clones used for tetraploid complementation. The ChIP-seq tracks show a complete absence of CTCF binding on the mutated CBS. The locations of the WT CBS are depicted with red and blue arrowheads (forward (F) and reverse (R) oriented respectively). Note that R1 and R2 in mESC display different binding enrichment as in limb data (**Fig. 1B**), denoting tissue-specific variation. **B.** Zoom in of the R1 genomic region. Note the absence of CTCF binding in mutants with deleted R1 CBS

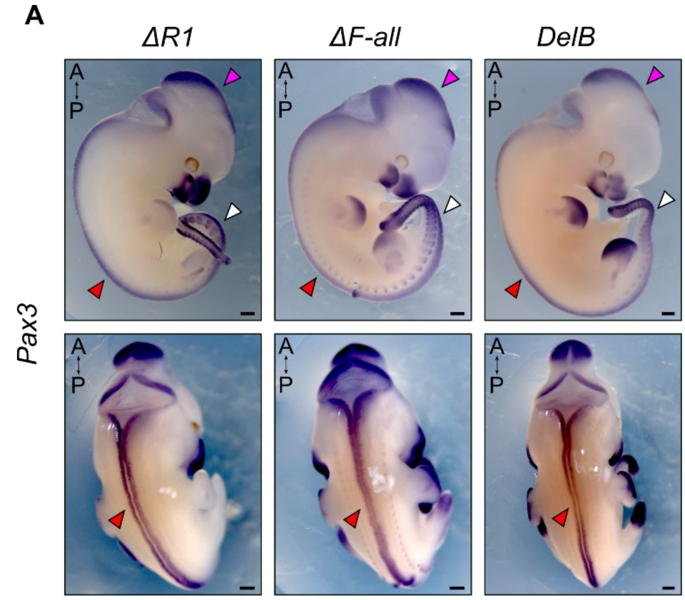


**Supp.Fig.6 Invariant *Pax3* expression pattern in different tissues.**

**A.** WISH shows *Pax3* expression in E11.5 mutants. Arrowheads indicate midbrain (purple), tail somites (white) and spinal cord (red). Note how the pattern of expression of *Pax3* is preserved in all tissues with the exception of the limb bud in the different CBS mutants. A, anterior; P, posterior. Scale bar: 500µm.

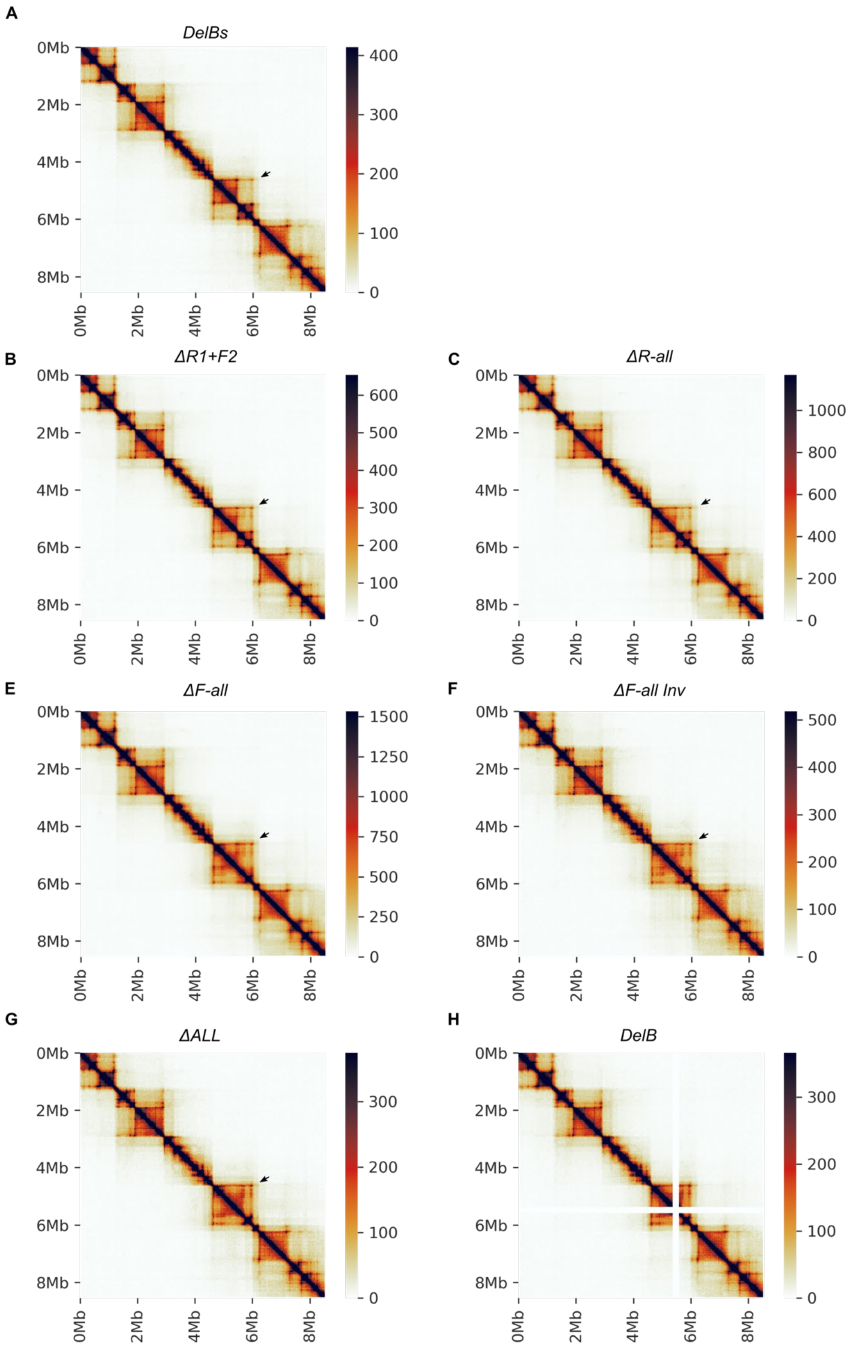


**Supp. Fig. 7. General view of the interaction matrices of embryonic distal limb cHi-C experiments**

**A-H.** Interaction matrices from embryonic limb cHi-C aligned to the custom genome representing the captured region (~8,3 Mb corresponding to the mm9 coordinates 71-81Mb excluding the *Epha4* intra-TAD deletion carried by the baseline *DelBs* mutant). The only exception is E (*ΔF-all Inv* mutant), where a modified version of the custom genome is used to account for the inversion of the boundary. Changes in the pattern of interactions of the different mutants are restricted to the progressive fusion of the *Epha4* and the *Pax3* TADs (black arrowheads).

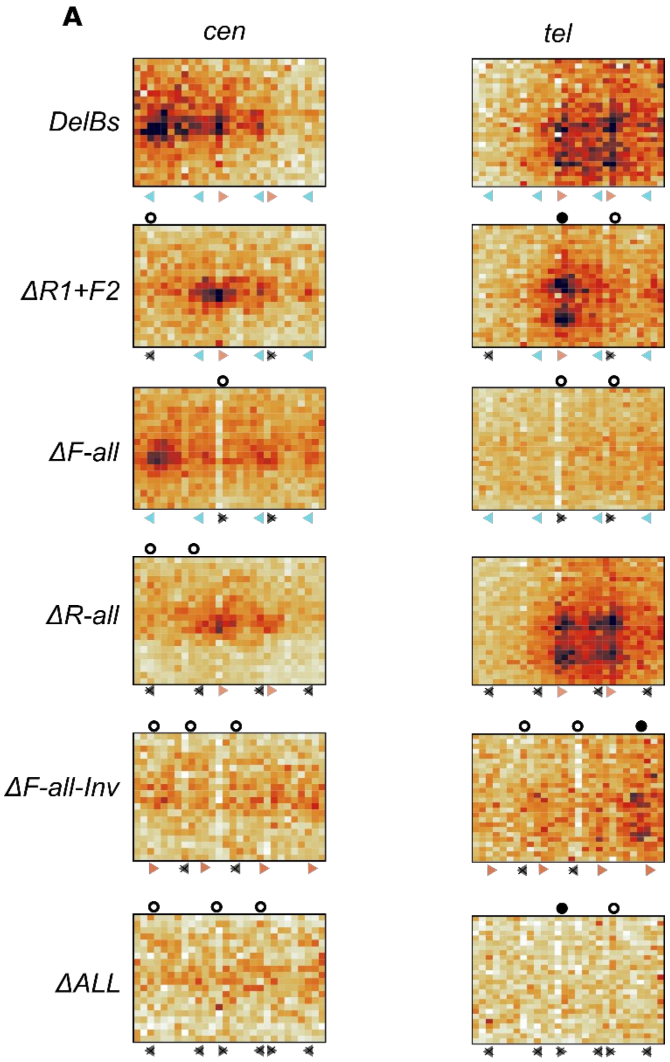


**Supp. Fig. 8. Non-convergent centromeric loops mediated by the F1 and F2 CBS.**

**A.** Close-up of the cHi-C interaction matrices showing the centromeric and telomeric loops established by the remaining CBS of the EP boundary in the different mutants. The coordinates shown are 4.55 Mb to 4.75 Mb for the centromeric loops and 5.8 Mb to 6.1 Mb for the telomeric. Genomic coordinates correspond to a custom *DelBs* genome for the captured region.

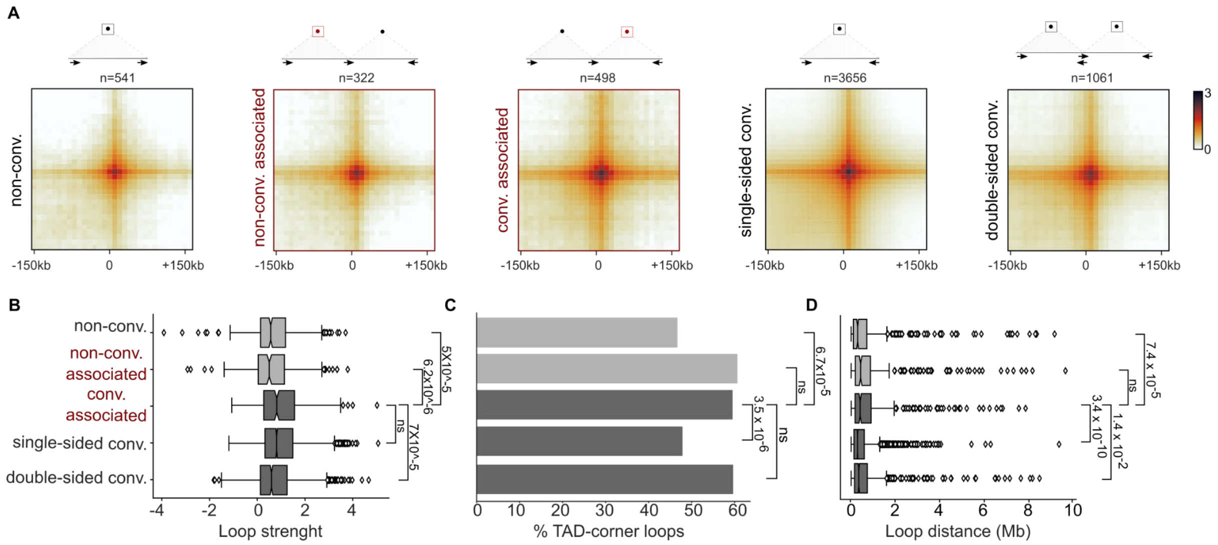


**Supp. Fig. 9. Paired convergent/non-convergent loops display longer distances between anchors and more association to TAD-corner loops than unidirectional convergent loops.**

**A.** Schematic (above) and loop aggregate plots (below) for all possible loop categories depending on the anchors: convergent (conv.) and non-convergent (non-conv.). Convergent loops can belong to the conv. associated if they share an anchor with a non-convergent loop in the opposite direction. If not, they can be either single-sided (if both their anchors establish loops in a single orientation) or double-sided. Non-convergent loops are further subdivided in non-conv. associated if they share an anchor with a convergent loop in the opposite direction or simply non-conv if they do not. Non-conv. associated and conv. associated loops are depicted in red because these categories associate to each other. **B.** Boxplots show the loop strength for the loop categories described in A. **C.** Barplots show the percentage of loops associated with putative TAD-corner loops for each of the categories. Significant differences between convergent associated loops and the rest of categories are highlighted with post-hoc $\chi 2$ p-values when appropriate. **D.** Boxplots represent the distribution of distances between anchors for the different loop categories. Significant post-hoc Mann-Whitney U p-values are shown in the appropriate comparisons between *conv. associated* loops and the rest of categories.

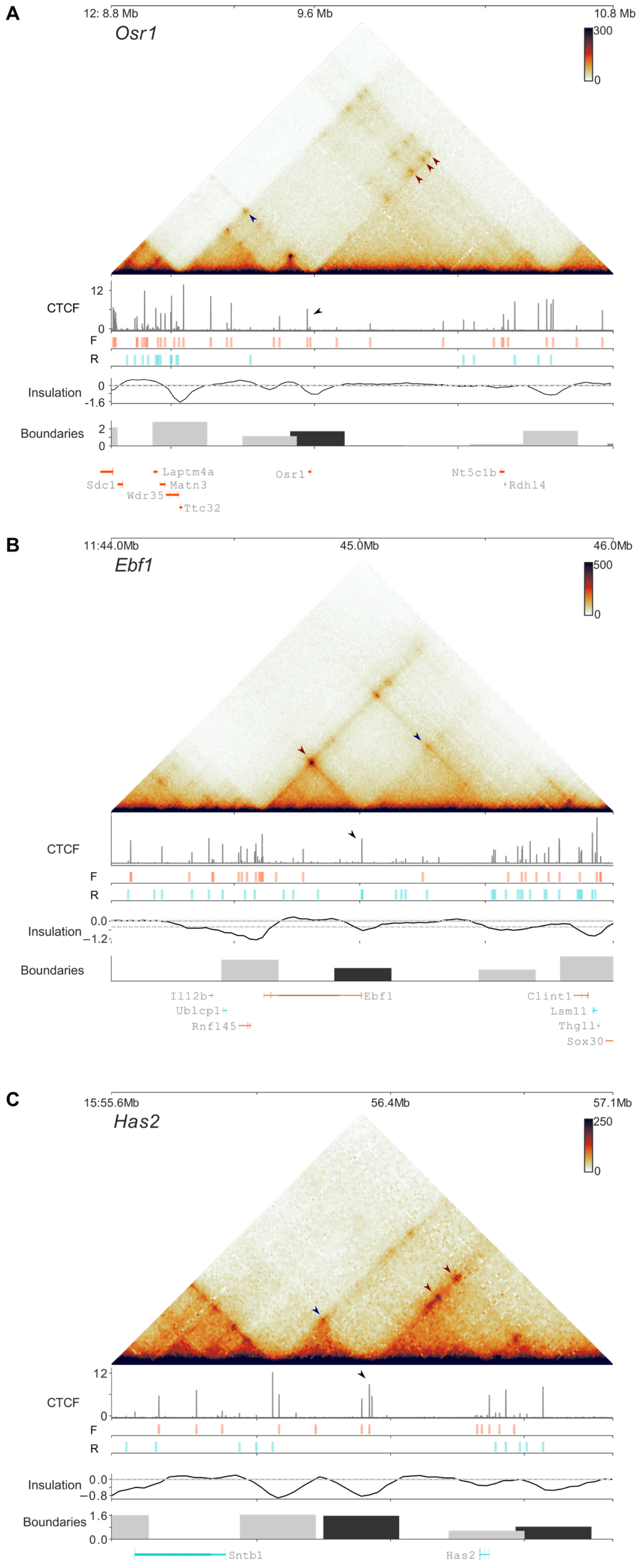
A
12: 8.8 Mb
9.6 Mb
10.8 Mb
Osr1
300
0
12
CTCF
0
F
R
Insulation
0
-1.6
Boundaries
2
0
Laptm4a
Osr1
Nt5c1b
Sdc1
Matn3
Rdh14
Wdr35
Ttc32
B
11:44.0Mb
45.0Mb
46.0Mb
Ebf1
500
0
CTCF
F
R
Insulation
0.0
−1.2
Boundaries
Il12b
Ebf1
Clint1
Ublcp1
Lsm11
Rnf145
Thg1l
Sox30
C
15:55.6Mb
56.4Mb
57.1Mb
Has2
250
0
12
CTCF
0
F
R
Insulation
0.0
−0.8
Boundaries
1.6
0.0
Sntb1
Has2

**Supp. Fig. 10. Loop anchors with strong single-oriented CTCF binding can constitute the source of weaker loops in the non-convergent direction.**

**A-C.** Three different examples of developmental gene loci with loop anchors that display unidirectional CBS and are engaged in both *convergent* and *non-convergent* loops. Hi-C interaction matrices and CTCF ChIP-seqs are from the mESC dataset in Bonev et al. 2017. CBS orientations are displayed in red and blue for positive and negative strands respectively and were calculated using FIMO (Grant et al. 2011, see methods). Dark red and dark blue arrowheads indicate *convergent* and associated *non-convergent* loops respectively. Insulation scores, boundaries and boundary scores were calculated with FAN-C (Kruse et al. 2020, see methods). Black boundary bars depict boundaries that do not contain divergent CBS pairs in their vicinity.

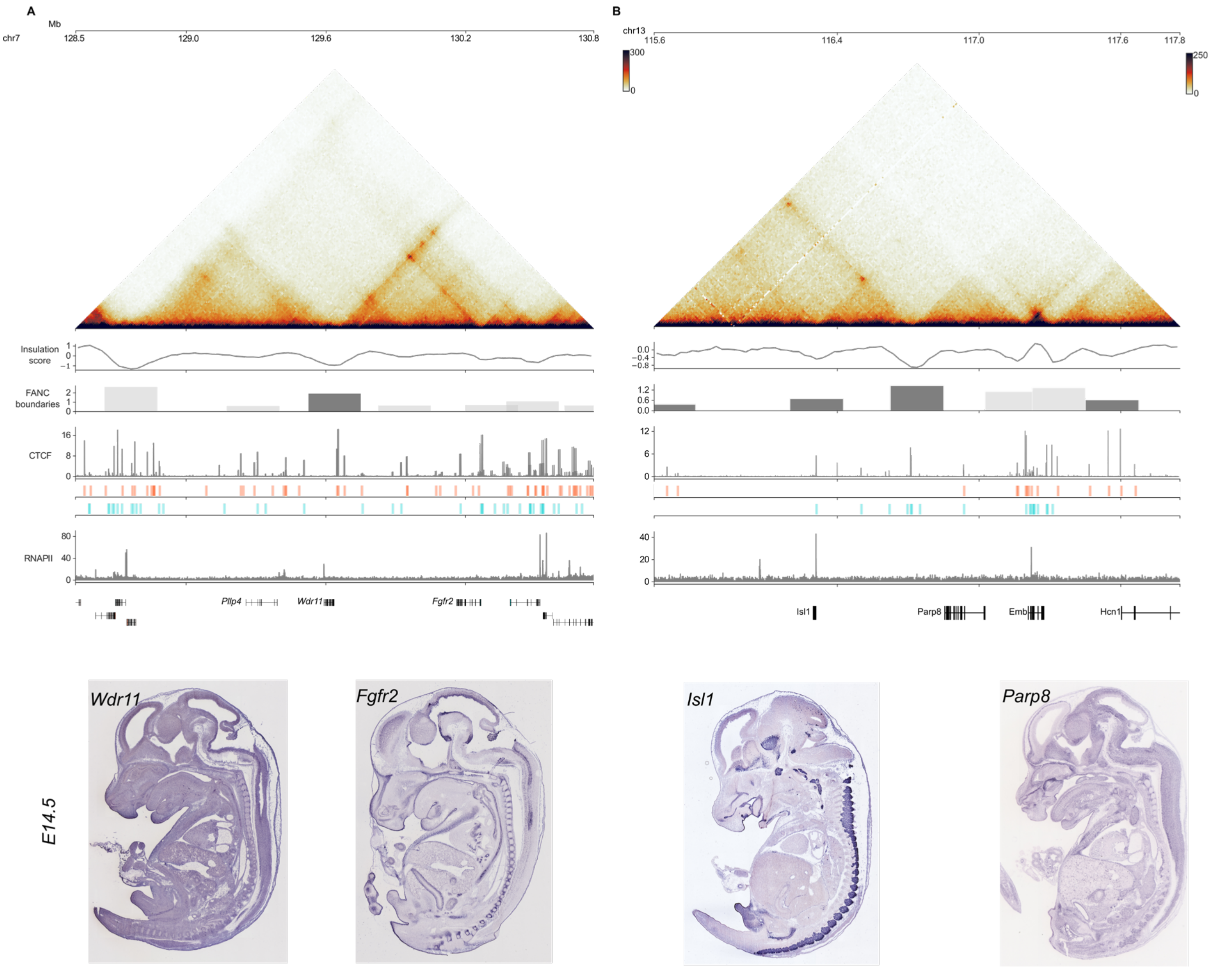


**Supp. Fig. 11. Boundary elements containing CBS in a single orientation can achieve comparable levels of insulation compared to boundaries containing divergent CBS.**

Two different examples of developmental loci, **(A)** *Fgfr2* and **(B)** *Isl1,* where boundary elements containing single-oriented CBS achieve boundary scores higher than one. Above, Hi-C and CTCF ChIP-seq experiments from mESC are shown (Bonev et al. 2017). CBS are depicted in red or blue for forward and reverse orientation respectively. Insulation scores, boundaries and boundary elements are calculated with FAN-C (Kruse et al. 2020, see methods). RNAPII ChIP-seq experiments in mESC (Sabari et al. 2018) do not show a particular enrichment in either of the boundaries. Below, E14.5 WISH from representative genes at either side of the boundary do not suggest co-regulation (obtained from GenePaint.org, Visel et al. 2004).

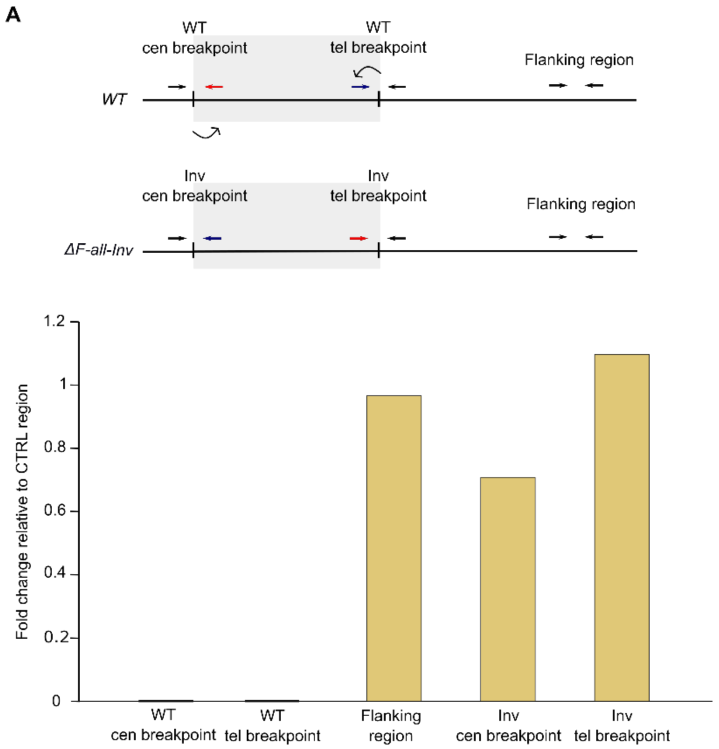


**Supp.Fig.12. *ΔF-all-Inv* genotyping.**

**A.** Schematics showing primer pairs positions (upper panel) and corresponding qPCR quantifying copy number of the inverted region in mutant mouse embryonic stem cells (lower panel). Bars represent the mean of three technical replicates. Values are normalized on a CTRL region on Chr13. Note how the *ΔF-all-Inv* mutant does not show any copy of the WT breakpoints. Cen, centromeric. Tel, telomeric

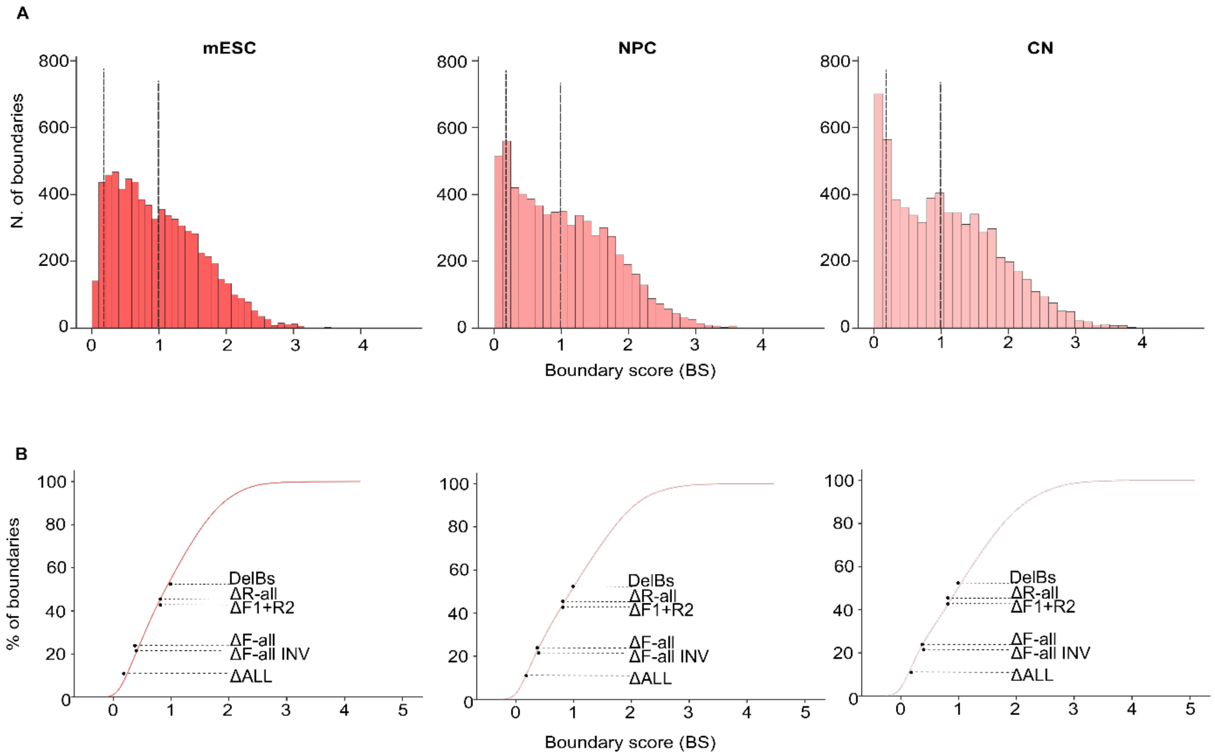


**Supp. Fig. 13. The boundary score (BS) of 40% of boundaries genome-wide could potentially allow regulatory inter-boundary interactions.**

**A.** Histogram representing the distribution of Boundary Scores genome-wide calculated from the mESC (left), neural progenitor cells (center) and cortical neurons (right) Hi-C datasets (Bonev et al. 2017). Many of them fall within the range of boundary scores of the EP boundary in our mutant series (demarcated by the vertical dashed lines). **B.** Cumulative distribution of the boundary scores from A. The boundary scores of the EP boundaries in each of our mutants is highlighted.
